# NuMA promotes constitutive heterochromatin compaction by stabilizing linker histone H1 on chromatin

**DOI:** 10.1101/2025.07.08.663620

**Authors:** Yao Wang, Wenxue Zhao, Jiahao Niu, Cuifang Liu, Xiaotian Wang, Weihong Yuan, Shanshan Ai, Wolfgang Baumeister, Guohong Li, Aibin He, Peng Xu, Cheng Li, Yujie Sun

## Abstract

Heterochromatin has been widely recognized to exert pivotal functions of silencing specific genes and maintenance of genome stability. However, the mechanisms underlying heterochromatin formation and maintenance remain to be fully elucidated. Here, we discovered that the critical mitotic regulator NuMA, as a nucleoskeleton protein, is required for constitutive heterochromatin organization at the level of nucleosomes in the interphase. Depletion of NuMA results in shortened nucleosome repeat length (NRL), dispersed nucleosome clutches and increased chromatin accessibility in heterochromatin regions. Afterwards, epigenetic maintenance and transcription repression in constitutive heterochromatin are disrupted upon NuMA-depletion, particularly the up-regulated transcription level of the non-coding long terminal repeat (LTR) elements, indicating the crucial roles of NuMA in cell differentiation and senescence. We revealed that such functions of NuMA rely on its interaction with linker histone H1, which stabilizes H1’s binding to chromatin and facilitates nucleosome stacking. We provided direct structural evidence of NuMA’s stabilization effect at the highest spatial resolution of nucleosomes through *in situ* cryo-ET. Notably, we found that NuMA oligomerizes into quasi-meshwork in nucleoplasm and highly co-localizes with H1 on the chromatin, providing the organization basis for NuMA as a nucleoskeleton protein in chromatin architecture regulation. Collectively, our findings illuminate the concerted effect of nucleoskeleton protein and linker histone on chromatin compaction at the level of nucleosomes, which unveil a new layer of mechanisms by which nucleoskeleton regulates heterochromatin formation and maintenance.

## Introduction

Heterochromatin refers to transcriptionally repressive chromatin domains essential for genome integrity and cellular identity. Classified by developmental dynamics and genomic distribution, facultative heterochromatin exhibits cell type or differentiation state-specific silencing patterns, whereas constitutive heterochromatin usually remains permanently condensed throughout the cell cycle and is enriched in pericentromeric regions, telomeres, and non-coding repetitive DNA elements. The assembly and maintenance of heterochromatin relies on sophisticated interplays among histone modifications (H3K27me3 for facultative heterochromatin and H3K9me2/H3K9me3 for constitutive chromatin) and various chromatin effectors, such as chromatin-remodeling complexes and heterochromatin protein 1 (HP1) family proteins^1–5^. Despite current progress, the molecular mechanisms underlying heterochromatin establishment and maintenance remains incompletely resolved, with a substantial number of regulatory factors awaiting discovery.

Emerging evidences implicate that nucleoskeleton participates in chromatin organization^6–9^ and heterochromatin structural regulation^10, 11^. Nucleoskeleton, also called nuclear matrix, was first termed in 1970s, and coined to describe the filamentous meshwork remained visible in nuclei subjected to high salt extraction and DNase digestion^12^. Notably, NuMA (Nuclear Mitotic Apparatus), a 238 kDa^13^ nucleoskeleton protein^14, 15^ displays dual-phase localization, spindle pole enrichment during mitosis and the nucleoplasm in interphase^16^ at a considerably high abundance (around 10^6^ copies *per* nucleus)^17–19^. While its mitotic roles in spindle assembly have been well-established^19, 20^, its functions in interphase remain enigmatic. Several clues suggest NuMA’s chromatin regulatory potential. Firstly, NuMA was proved to bind DNA *in vitro via* its C-terminal domain^21–23^. Secondly, previous studies showed that NuMA contributes to proper chromatin decompaction and nuclear shape by directly associating with DNA at mitotic exit^22, 23^, and down-regulation of NuMA accelerates apoptotic destruction of nuclei in MCF-7 cells^24^ and perturbs tissue-specific epigenetic modifications in breast epithelial cultures^25^. Besides, over-expressed NuMA forms meshwork-like lattice structure in HeLa nuclei, and purified NuMA self-assembles into ‘multi-armed’ oligomers *in vitro*^13, 26^, suggesting its scaffold functions. Significantly, NuMA was reported to exhibit interaction with histone H1.2 in the yeast two-hybrid system^26^. Considering linker histone H1’s central roles in binding the linker DNA of nucleosome core particles (NCPs) to stabilize nucleosome structure and condense chromatin^27^, there is great potential for NuMA to participate in heterochromatin regulation.

In this study, we thoroughly explored the functions and unveiled the molecular mechanisms of NuMA in interphase chromatin organization by combining super-resolution imaging, high-throughput sequencing, cryo-ET and a variety of biophysical and biochemical assays, and proposed that NuMA promotes constitutive heterochromatin compaction and transcriptional repression of LTR through stabilizing histone H1 on chromatin.

## Results

### NuMA regulates global 3D genome organization in the interphase

To investigate NuMA’s functions specifically in the interphase, we constructed an auxin-inducible degradation (AID) system^28^ in the HCT116 cell line given the lethality of NuMA knock-out in mice as demonstrated in previous research^29^. Through adding auxin at the end of second round synchronization (See Methods for details), the protein level of NuMA within the interphase time range was rapidly reduced (Figure 1A). Based on the AID system, we explored the effect of NuMA-depletion on the nucleus and genome organization during the interphase.

**Figure 1.**
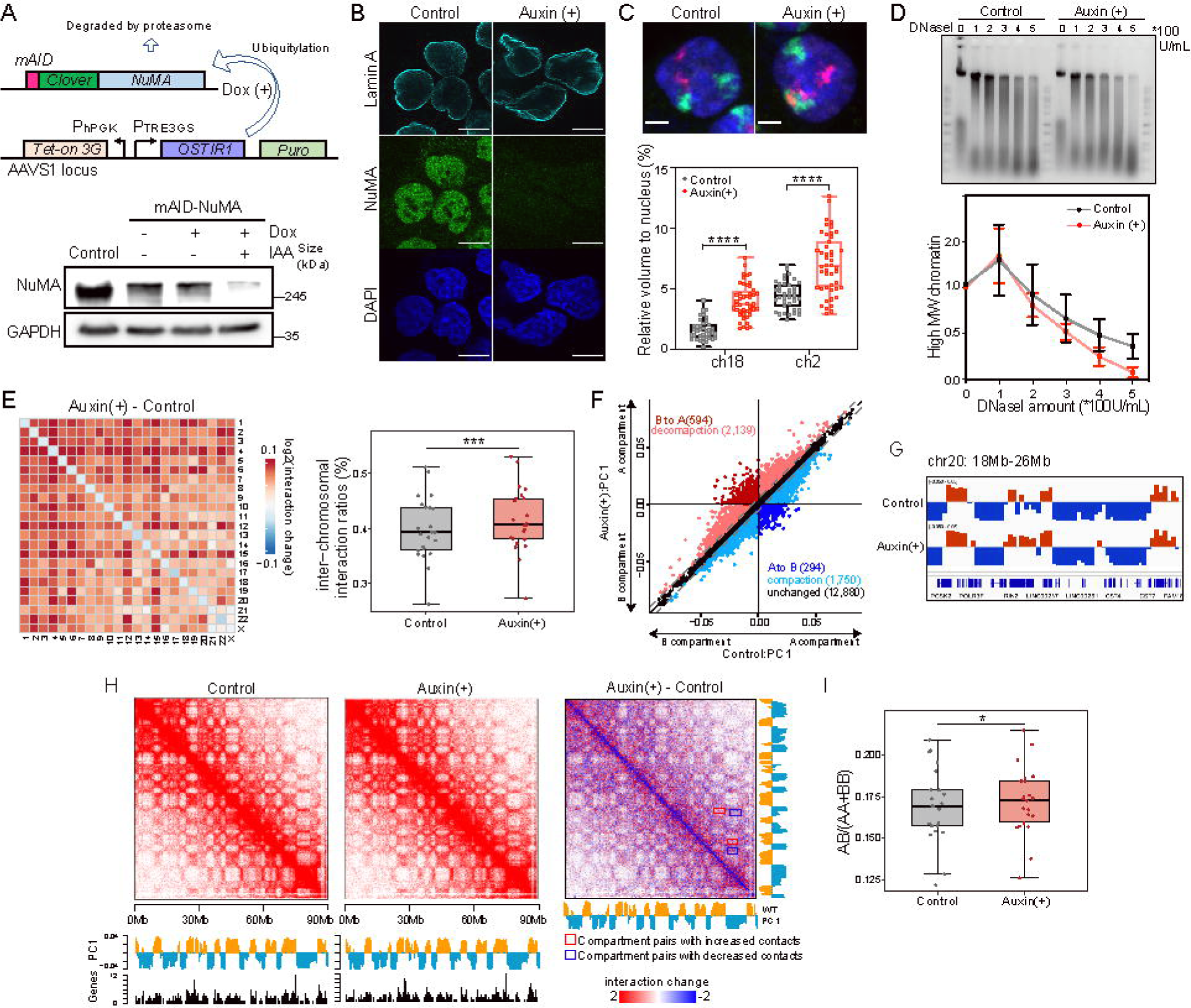
NuMA regulates global 3D genome organization. A. Schematic of auxin-inducible degron system (upper) and NuMA-depletion induced by auxin in unmodified HCT116 cells as control and HCT116-mAID-NuMA cells detected by western blotting (lower). B. IF imaging of Lamin A in untreated HCT116-mAID-NuMA cells as control and NuMA-depleted HCT116-mAID-NuMA cells induced by auxin. Scale bar, 10 μm. C. FISH imaging (upper) and quantification (lower) of the volumes occupied by chromosome 2 (green) and 18 (red) relative to the nucleus volume in untreated HCT116-mAID-NuMA cells as control and NuMA-depleted HCT116-mAID-NuMA cells induced by auxin. Error bars represent SD (n≥50). ****p < 0.0005, Mann-Whitney test. Scale bar, 5 μm. D. Global chromatin accessibility upon NuMA-depletion detected by DNaseI digestion assay, with untreated HCT116-mAID-NuMA cells as control. Agarose gel image of genomic DNA digested by DNaseI at different concentrations (left), and percentages of high-molecular-weight (MW) genomic DNA (>5 kb) (right). The gel is representative of three independent experiments. Error bars represent SD (n=3). E. Heat map acquired by subtracting the inter-chromosomal ratios in untreated HCT116-mAID-NuMA cells as control and NuMA-depleted HCT116-mAID-NuMA cells induced by auxin (left). Trans-interaction ratios of each chromosome in control and NuMA-depletion induced by auxin in HCT116-mAID-NuMA cells (right). ***P < 0.001, paired t-test. F. Scatterplot showing compartment switches in untreated HCT116-mAID-NuMA cells as control and NuMA-depleted HCT116-mAID-NuMA cells induced by auxin. PC1 was calculated for each 150 kb genomic segment to define A and B compartments and identify compartment switching, decompaction and compaction upon NuMA-depletion. G. A representative region showing the compartment switch from B to A upon NuMA-depletion. H. Heat map (resolution: 150 kb) acquired by subtracting compartment contacts in untreated HCT116-mAID-NuMA cells as control and NuMA-depleted HCT116-mAID-NuMA cells induced by auxin, and differential matrices of NuMA-depleted minus control cells. Below the heatmaps are PC1 values and gene density plots. I. Compartment A and B interaction ratios in untreated HCT116-mAID-NuMA cells as control and NuMA-depleted HCT116-mAID-NuMA cells induced by auxin. *P < 0.05, wilcoxon test for paired samples.

Firstly, immunofluorescence (IF) of the nuclear envelope labeled by lamin A showed different degrees of shrinkage after NuMA-depletion (Figure 1B), suggesting that NuMA plays a role in nuclear mechanical maintenance, corroborating earlier findings^23^. Next, fluorescent *in situ* hybridization (FISH) of chromosome 2 and chromosome 18 which respectively represent chromosomes with relatively higher and lower gene content showed that NuMA-depletion led to significant increase of chromosome volume (Figure 1C). This prompted us to explore whether the enlargement of chromosome is accompanied by alterations in chromosome compaction. To answer this question, we performed DNaseI digestion assay to assess the global chromatin accessibility. The results showed that the genome became more sensitive to DNaseI digestion after NuMA-depletion (Figure 1D), indicating a more accessible and less compacted chromatin status. Collectively, these results suggested that NuMA is involved in nuclear organization and promotes chromatin compaction in the interphase. To further verify these findings, we also constructed the AID system in the U2OS cell line (Figure S1A) and observed similar phenotypes upon NuMA-depletion, including shrinkage of nuclear envelope (Figure S1B), enlarged chromosomes volume (Figure S1C), and increased chromatin accessibility (Figure S1D).

Based on the observed critical role of NuMA in chromatin spatial organization, we performed high-throughput chromosome conformation capture (Hi-C) assay^30^ in control and NuMA-depleted HCT116 cells to measure the hierarchical genome architecture alterations (S1E-G). At the whole genome level, while global chromosomal contact frequency did not exhibit significant changes (Figure S1F and S1G), the inter-chromosomal interactions increased (Figure 1E), in line with the enlarged chromosome volume revealed by FISH (Figure 1C) and enhanced chromatin accessibility examined by the DNaseI digestion assay (Figure 1D). Compartment-level analysis demonstrated prominent decompaction in compartment B, with predominant B-to-A transitions over A-to-B (Fig. 1F and 1G), suggesting that NuMA is important for chromatin compartmentalization and chromatin compaction maintenance in B compartments. The significantly elevated ratio of inter-compartment interactions also suggests a weaker segregation of different chromosomal compartments (Figure 1H and 1I). Taken together, these results indicated that NuMA is required for 3D genome organization at both the chromosome and compartment scales, whose depletion induces global chromatin decompaction, particularly for B compartments.

### NuMA regulates heterochromatin compaction by maintaining nucleosome stacking

To quantitatively assess global chromatin decompaction induced by NuMA-depletion, we performed transposase-accessible chromatin using sequencing (ATAC-seq)^31^ assay in control and NuMA-depleted HCT116 cells to map the alterations in chromatin accessibility across the genome. We constructed a chromHMM^32^ chromatin state map which classified the whole genome into 15 distinct categories for analyzing NuMA’s impact on chromatin states throughout the genome (Figure 2A). We compared the chromatin accessibility of control and NuMA-depleted cells reflected by ATAC-seq fragment length and density in the 15 categories of chromatin, and calculated the nucleosome-repeat length (NRL) as previously described^33^. According to the changes in chromatin accessibility and NRL in response to NuMA-depletion, the 15 categories of chromatin exhibited three distinct patterns, which we defined as Type 1, 2 and 3 chromatin (Figure 2B, S2A). Type 1 chromatin demonstrated the most significant increase of ATAC-seq reads frequency and reduction of NRL while Type 2 chromatin exhibits slight increase in chromatin accessibility, and neither of Type 2 or Type 3 chromatin showed obvious changes in NRL after NuMA-depletion (Figure 2B, S2A). Significantly, Type 1 chromatin is predominantly comprised of heterochromatic domains enriched with H3K9me3, H3K27me3, and unannotated post-translational modifications, while Type 2 and 3 chromatin encompasses transcription-active and transcription-regulatory regions. Analysis of Hi-C data according to the categorization of Type 1, 2 and 3 chromatin also showed that compartment B was significantly concentrated in Type 1 chromatin (Figure S2B), further verifying that NuMA-depletion mainly affects genome 3D organization in heterochromatin (Figure 1F, G). Furthermore, in regions undergoing compartment A/B switch (including A to B and B to A), Type 1 chromatin also accounted for the highest proportion (Figure S2C). Focusing specifically on compartment A/B switches within Type 1 chromatin, we found that the compartment B to A switch and chromatin de-compaction also accounted for the highest proportion (Figure S2D). Collectively, the integrated analysis of ATAC-seq and Hi-C data indicated that NuMA-depletion leads to a more accessible chromatin state and shortening of NRL in heterochromatin regions, suggesting that NuMA regulates heterochromatin compaction and nucleosome spacing.

**Figure 2.**
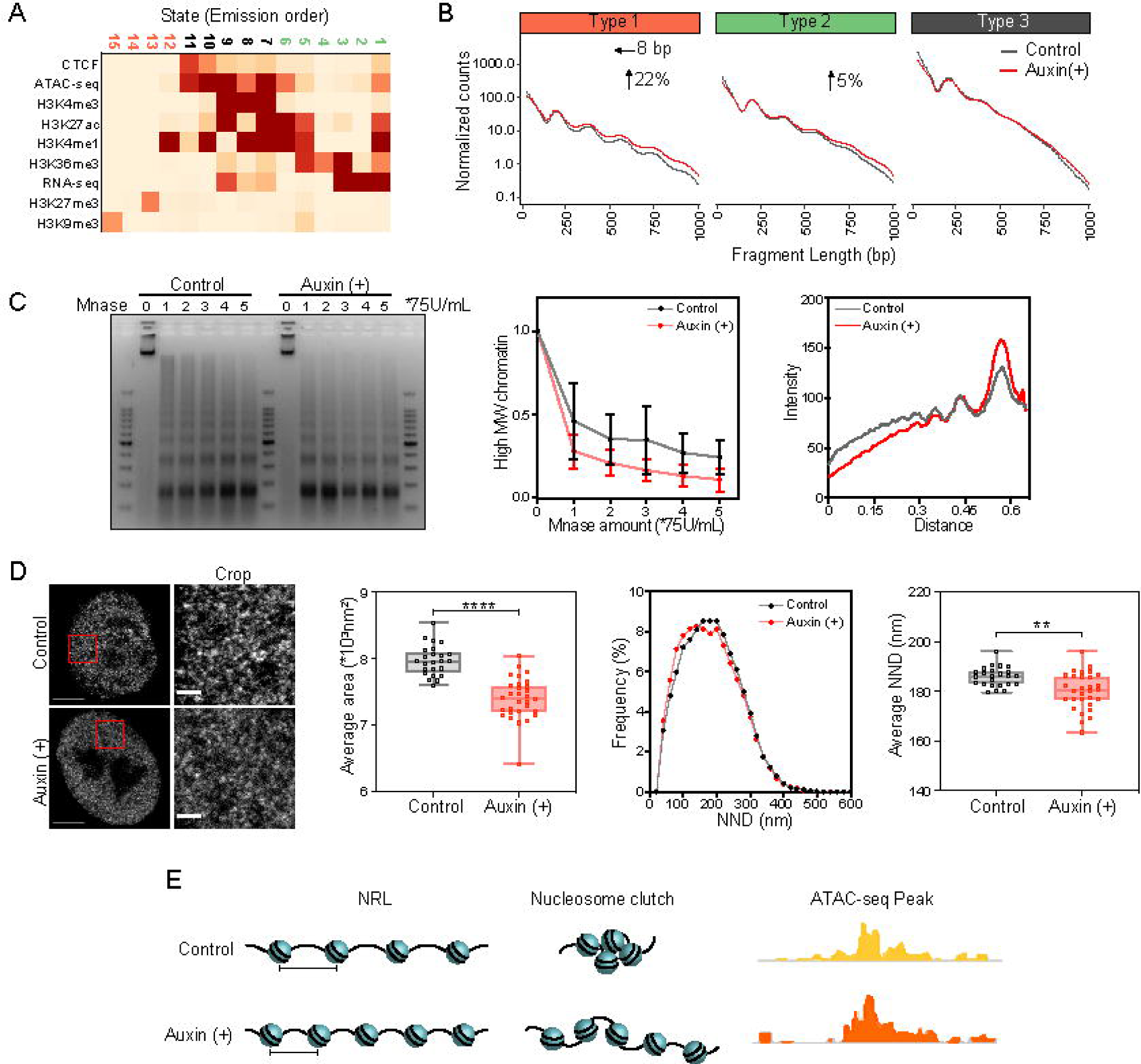
NuMA regulates heterochromatin compaction by maintaining nucleosome stacking. A. A 15-state chromatin model established in HCT116 cells using ChromHMM. The color of each square represents the enrichment degree of chromatin feature. B. ATAC-seq fragment length and density of untreated HCT116-mAID-NuMA cells as control and NuMA-depleted HCT116-mAID-NuMA cells induced by auxin classified by chromatin types (Type 1, 2, and 3). Changes in nucleosome repeat length (NRL) and chromatin accessibility are shown for each chromatin state. The change in accessibility and NRL are shown as determined by NRLfinder. C. Chromatin accessibility upon NuMA-depletion in HCT116-mAID-NuMA cells as detected by Mnase digestion assay. Gel image of genomic DNA digested by Mnase at different concentrations (left), percentages of high MW genomic DNA (>5 kb) (middle), and an example of nucleosome ladders at the well 2 is shown (right). The gel is representative of three independent experiments. Error bars represent SD (n=3). D. STED imaging of H2B (left) and quantification (right) of nucleosome clutches in HCT116-mAID-NuMA cells, including the average area, frequency of nearest neighbor distance (NND) and average NND. Error bars represent SD (n=20). ****p < 0.0005, **p < 0.05, Mann-Whitney test. Scale bar, 5 μm (left) and 1 μm (right). E. A cartoon model depicting the changes in NRL and chromatin accessibility upon NuMA-depletion.

We also conducted Mnase digestion assay and super-resolution imaging of nucleosome clutches^34^ to confirm the effect of NuMA-depletion on nucleosome spacing and stacking. The results showed that after NuMA-depletion in HCT116 and U2OS cells, chromatin’s sensitivity to Mnase was increased, which confirmed the shortening of NRL and increase of chromatin accessibility (Figure 2C, S2E). Labeling of H2B by immunofluorescence and imaging of nucleosomes *in situ* by Stimulated Emission Depletion (STED) imaging revealed that NuMA-depletion had a dispersing effect on nucleosome clutches, evident in reduced average area and decreased nearest neighbor distance (NND) between them (Figure 2D, S2F-H), suggesting NuMA’s effect on spatial arrangement of nucleosomes. As reported in the previous research, larger and denser clutches containing more nucleosomes formed “closed” heterochromatin, while smaller and less dense clutches formed “open” chromatin^34^ which aligns with the observation that NuMA mainly affect the accessibility of Type 1 chromatin (Figure 2B, S2B-D).

Collectively, these data demonstrated that NuMA-depletion disrupts genome architecture at the nucleosome level characterized by decreased NRL and reduced density of nucleosome spacing. These structural alterations provide us mechanistic evidence that NuMA governs heterochromatin compaction through maintaining nucleosome stacking (Figure 2E).

### NuMA stabilizes linker histone H1’s binding to chromatin and facilitates nucleosome stacking

As the most abundant chromatin-binding protein in eukaryotes, linker histone H1 binds to the entry and exit sites of DNA on the nucleosome core particles (NPCs) to stabilize nucleosome structure and compact chromatin^27, 35, 36^. H1 is one of the direct regulatory factors of NRL, and there is a robust linear relationship between H1 stoichiometry and NRL^27, 36, 37^. In 1998, Gueth-Hallonet and his colleagues reported that NuMA exhibits interaction with histone H1.2 in the yeast two-hybrid system^26^. Therefore, we hypothesized that the impact of NuMA-depletion on genome architecture, especially at the nucleosome level, is closely related to its interaction with H1.

The correlation between NuMA and H1 was substantiated by the co-localization of mScarlet-fused NuMA and eGFP-fused H1.1 revealed by live-cell imaging (Figure 3A). Moreover, we found that among the N-terminus (hereinafter abbreviated as NuMA-N), C-terminus (hereinafter abbreviated as NuMA-C) and 200 nm-long coiled-coil domain of NuMA (Figure S3A), only NuMA-C showed co-localization with H1.1 and six other H1 variants in somatic cells^27^ (Figure S3B), suggesting that NuMA interacts with H1 through its C-terminus. This finding was also verified by co-IP of endogenous NuMA with H1 (Figure S3C, S3D), overexpressed NuMA with H1 (Figure 3B, S3E), and overexpressed NuMA-C with H1 (Figure S3F). Besides, we mapped the interaction regions of NuMA binding to H1 through charactering the spatial localization by live-cell imaging and correlation by co-IP between overexpressed NuMA-C and H1. We identified two accurate regions of NuMA-C binding to H1, including region aa 1778-1942 and aa 2004-2115 (Figure 3H, S3G, S3H), which we named as H1BD1 (H1 binding domain 1) and H1BD2 (H1 binding domain 2), respectively. Through *in vitro* GST-pull down experiments, we also proved that NuMA-C directly binds H1 *via* H1BD1 and H1BD2 (Figure S3I, S3J).

**Figure 3.**
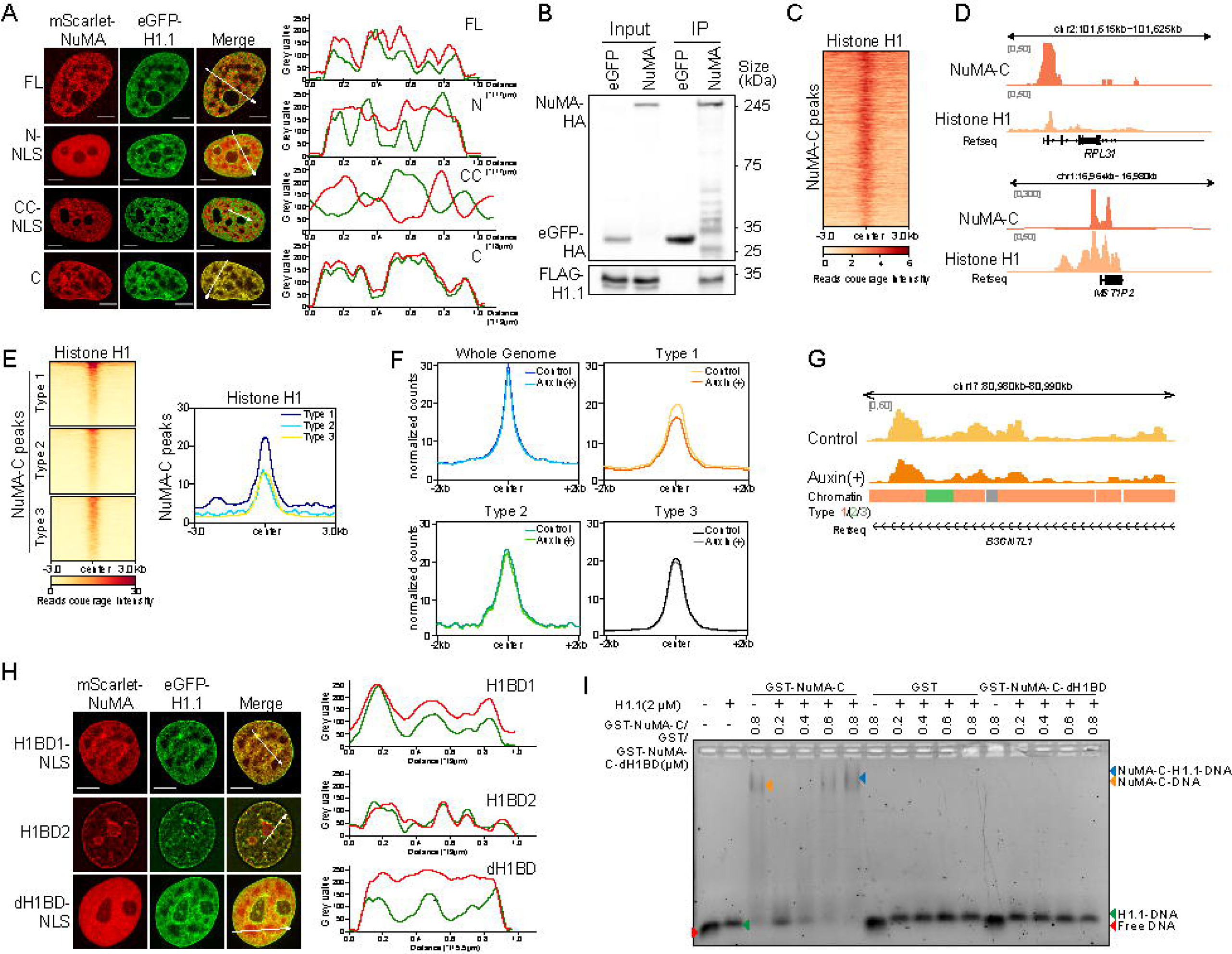
NuMA promotes linker histone H1’s binding to chromatin. A. Representative live-cell image of over-expressed NuMA truncations and H1.1 in HeLa cells (left). Plots of the red and green pixel intensities along the white arrow in the left panel (right). Scale bar, 5 μm. B. Immunoblots showing the immunoprecipitation of HA-tagged NuMA and FLAG-tagged H1.1 over-expressed in 293T cells. C. Heatmap profiling of H1’s CUT&Tag signal at NuMA-C’s regions, and the peaks were aligned using the center of the peaks. D. Representative examples of co-localization of NuMA-C and H1’s peaks. E. Heatmap profiling of H1’s CUT&Tag signal at NuMA-C’s regions in each chromatin type, and the peaks were aligned (left) using the center of the peaks and summed (right). F. CUT&Tag peaks of H1 in the whole genome and each chromatin type in untreated and NuMA-depleted HCT116-mAID-NuMA cells, and the peaks were aligned using the center of the peaks. G. Representative examples of H1 binding changes in each chromatin type. H. Representative live-cell image of over-expressed NuMA-C’s truncations and H1.1 in HeLa cells (left). Plots of the red and green pixel intensities along the white arrow in the left panel (right). Scale bar, 5 μm. I. NuMA-C and H1.1 bind DNA together in vitro shown by EMSA analysis.

NuMA was reported to bind DNA directly *in vitro* through its evolutionarily conserved last 58 amino acids in the C-terminus^21–23^. To profile NuMA’s *in situ* DNA binding sites across the whole genome, we performed cleavage under target & tagmentation (CUT&Tag)^38^ experiments using an antibody targeted to NuMA-C. The results showed that NuMA indeed exhibited specific DNA binding sites on the genome and highly co-localized with the peaks of H1 (Figure 3C and 3D). Interestingly, we observed the highest occupancy of NuMA in Type 1 chromatin (Figure S3K), and the degree of colocalization of NuMA and H1 was also the highest in Type 1 chromatin (Figure 3E). These data are consistent with the observation that NuMA-depletion mostly impacts the architecture of heterochromatin (Figure 2A and 2B).

Intrigued by the dual binding capacity of NuMA with H1 and DNA, we questioned whether NuMA can affect H1’s binding to chromatin and thereby regulate nucleosome stacking. We performed CUT&Tag experiments of H1 in control and NuMA-depleted HCT116-mAID-NuMA cells, and found that the coverage of H1’s binding sites and counts of peaks were both significantly reduced in Type 1 chromatin, in clear contrast to those in Type 2 and Type 3 chromatin (Figure 3F and 3G), which is consistent with NuMA’s occupancy and NuMA’s co-localization with H1 on genome (Figure 3E). To validate the ability of NuMA promoting H1’s binding on DNA *in vitro*, we performed electrophoretic mobility shift assay (EMSA), and found that in the presence of NuMA-C, H1’s DNA-binding capacity was significantly strengthened. In contrast, the truncation of NuMA which deleted its two H1BDs was unable to increase the binding of H1 to DNA (Figure 3I and S3L). Taken together, we concluded that NuMA stabilizes linker histone H1’s binding to chromatin through its interaction with H1.

Extensive studies have established linker histone H1’s nucleosome-stabilizing role in chromatin condensation through linker DNA binding. Besides of *in vitro* evidences^39, 40^, H1’s preferential enrichment in heterochromatin and large nucleosome clutches were also wildly recognized^34^. Therefore, we proposed that NuMA’s stabilization effect on H1’s binding to DNA is the mechanism by which NuMA regulates chromatin architecture at the nucleosome level.

To confirm this hypothesis, *in vitro* pull-down assay was applied to evaluate NuMA’s binding affinity to single nucleosomes. Immunoblotting results revealed that NuMA-C could bind to individual nucleosomes *in vitro*, and nucleosomes containing histone H1 pulled down significantly more NuMA-C (Figure 4A), suggesting the binding preference of NuMA to H1-associated nucleosomes.Then we performed atomic force microscopy (AFM) to characterize NuMA’s effect on the conformation of nucleosome array (NA). As shown in Figure 4B, the 12× NA adopted stretched beads-on-string conformation, and the addition of H1 led to a condensed structure. Afterwards, the addition of NuMA-C caused compacted 12× NA to form larger condensates in the presence of H1. NuMA’s enhancement on H1’s compaction effect on 12× NA was also confirmed by the sucrose density-gradient centrifugation result (Figure 4C). Noteworthily, we found that without the presence of H1, NuMA-C showed no significant effects on either the conformation of 12× NA (Figure 4B) or the sedimentation coefficient (Figure 4C), suggesting that NuMA’s regulatory effect on chromatin architecture is dependent on its interaction with H1.

**Figure 4.**
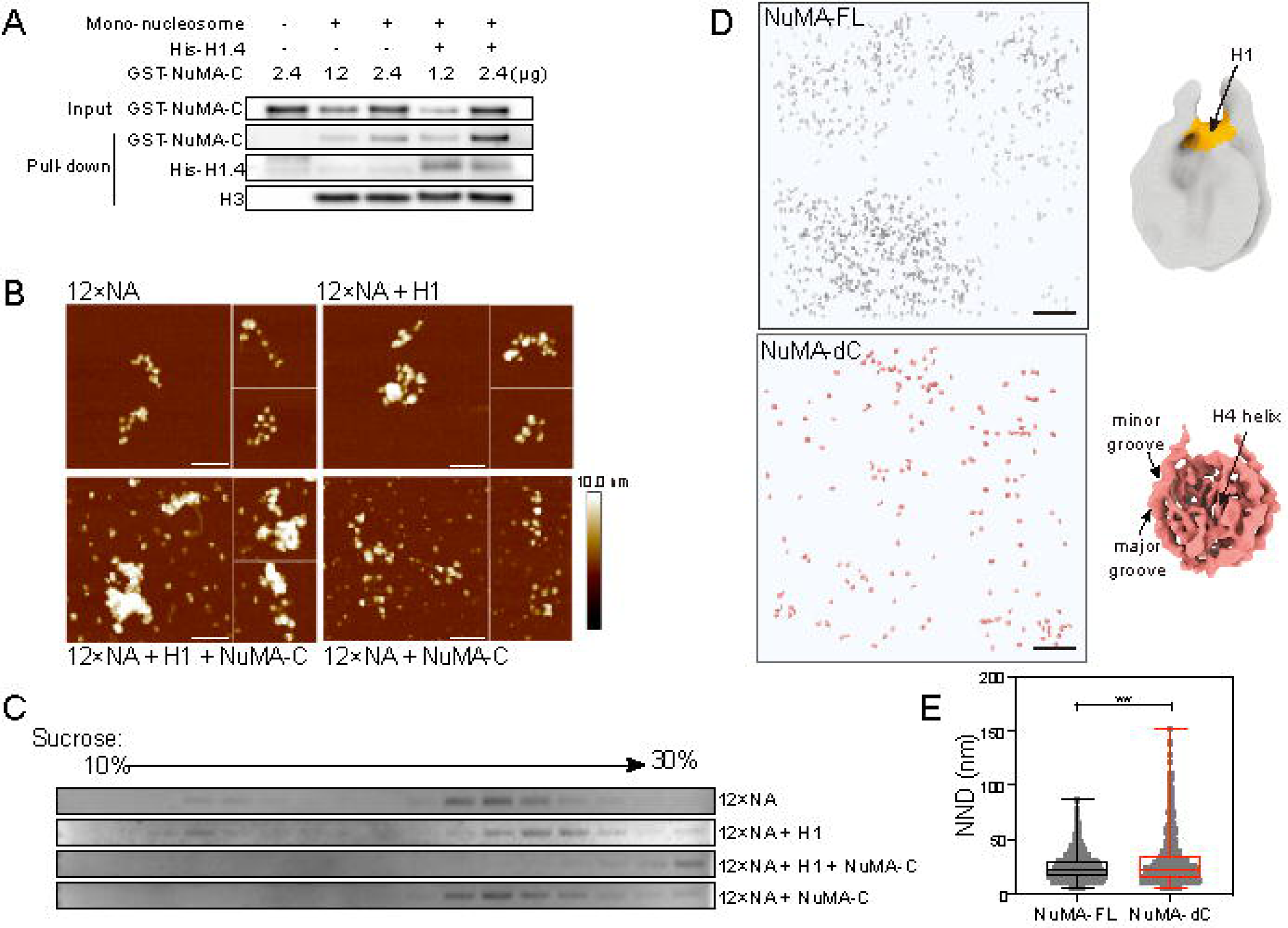
NuMA facilitates nucleosome stacking by stabilizing H1. A. The binding preference of NuMA-C to nucleosomes with H1 as detected by mono-nucleosome pull-down experiment and western blotting. B. AFM images for the reconstituted 12× nucleosome arrays with H1.4 and NuMA-C. Scale bar, 100 nm. C. Sucrose density gradient centrifugation result of 12× nucleosome array with H1.4 and NuMA-C. DNA displayed by EB staining. D. Upper left, 3D rendering from tomograms of U2OS cells overexpressing full-length NuMA (NuMA-FL) Lower left, 3D rendering from tomograms truncation deleting the C-terminal domain (NuMA-dC). Scale bar, 100 nm. Upper right, *in situ* nucleosome subtomogram averaging electron density map from cells overexpressing NuMA-FL. Lower right, *in situ* nucleosome subtomogram averaging electron density map from cells overexpressing NuMA-dC. H1 linker region labeled orange. Features labeled with black arrows. E. *In situ* nucleosome NND distribution. **p < 0.05, Mann-Whitney test.

To elucidate NuMA’s facilitation effect on nucleosome stacking under conditions close to native states, we conducted advanced cryo-focused ion beam (FIB) milling combined with cryo-electron tomography (cryo-ET) in U2OS cells overexpressing full-length NuMA (hereinafter abbreviated as NuMA-FL) and a truncated variant of NuMA deleting the C-terminus (hereinafter abbreviated as NuMA-dC, Figure S3A). Tilt-series analysis revealed that nucleosomes were more compactly arranged with a significantly reduced nearest neighbor distance (NND) in cells overexpressing NuMA-FL compared to NuMA-dC (Figure 4D, 4E, S4A). This difference observed between the overexpression of NuMA-FL and NuMA-dC under identical experimental conditions precisely underscored the pivotal role of NuMA-C in nucleosome stacking maintenance. At the same time, subtomogram averaging resolved the conformation of NCPs with nanometer resolution at 9.9 Å from NuMA-dC tomograms, allowing proper fitting of histone helices into the electron density map (Figure S4B, S4D, S4E, 4D). In parallel, a chromatosome conformation was resolved at 17.8 Å resolution from NuMA-FL tomograms using the exactly same workflow as NuMA-dC (Figure S4C, S4D, S4F, 4D). Notably, a distinct density was resolved at the DNA entry/exit site, which could be accurately fitted with the H1 globular domain (Figure 4D, S4F). In contrast, no extra density was detected at the H1 linker region throughout the entire processing workflow from NuMA-dC (Figure S4E), highlighting the stabilization influence of NuMA on H1’s binding to chromatin. Collectively, these results demonstrate that NuMA facilitates nucleosome stacking through stabilizing the interaction between H1 and nucleosome core.

### NuMA oligomerizes into quasi-meshwork organization through its C-termini *in vivo*

Previous studies reported that purified NuMA self-assembles into ‘multi-armed’ oligomers through its C-terminus *in vitro* and over-expressed NuMA forms lattice structure in HeLa nuclei^13, 26^, suggesting its structural and supportive functions. Since we have proved NuMA’s interaction with H1 and regulatory effect on it, we performed super-resolution imaging to delineate the distribution pattern of endogenic NuMA in the nucleoplasm to seek the structural basis of NuMA for its genome organization functions.

To achieve this, we co-labeled the two ends of NuMA with antibodies specifically targeting to NuMA-N and NuMA-C, respectively (Figure 5A). Due to its considerably high abundance^17–19^, both confocal imaging and STED super-resolution imaging were insufficient to resolve individual NuMA molecules (Figure S5A). We therefore performed expansion microscopy as previously described^41, 42^, which enabled characterization of Atto647N-labeled NuMA-N and Alexa594-labeled NuMA-C in U2OS cells (Figure 5B). In order to unveil the organization pattern of NuMA molecules, we linked the foci of NuMA-N and NuMA-C within its molecular length, *i.e.* 250 nm^13^, and calculated the stoichiometry and NND between NuMA-N and NuMA-C (Figure 5B). As seen in the cropped view, a NuMA-C focus was encircled by 2 ∼ 4 NuMA-N foci, which is similar with the pattern observed *in vitro*^13^. The average NuMA-N to NuMA-C ratio was 2.808, and the mean NND from NuMA-C to NuMA-N clusters was 107.22 nm in U2OS cells (Figure 5C). These results suggested that NuMA forms a quasi-meshwork organization *via* oligomerization of NuMA-C.

**Figure 5.**
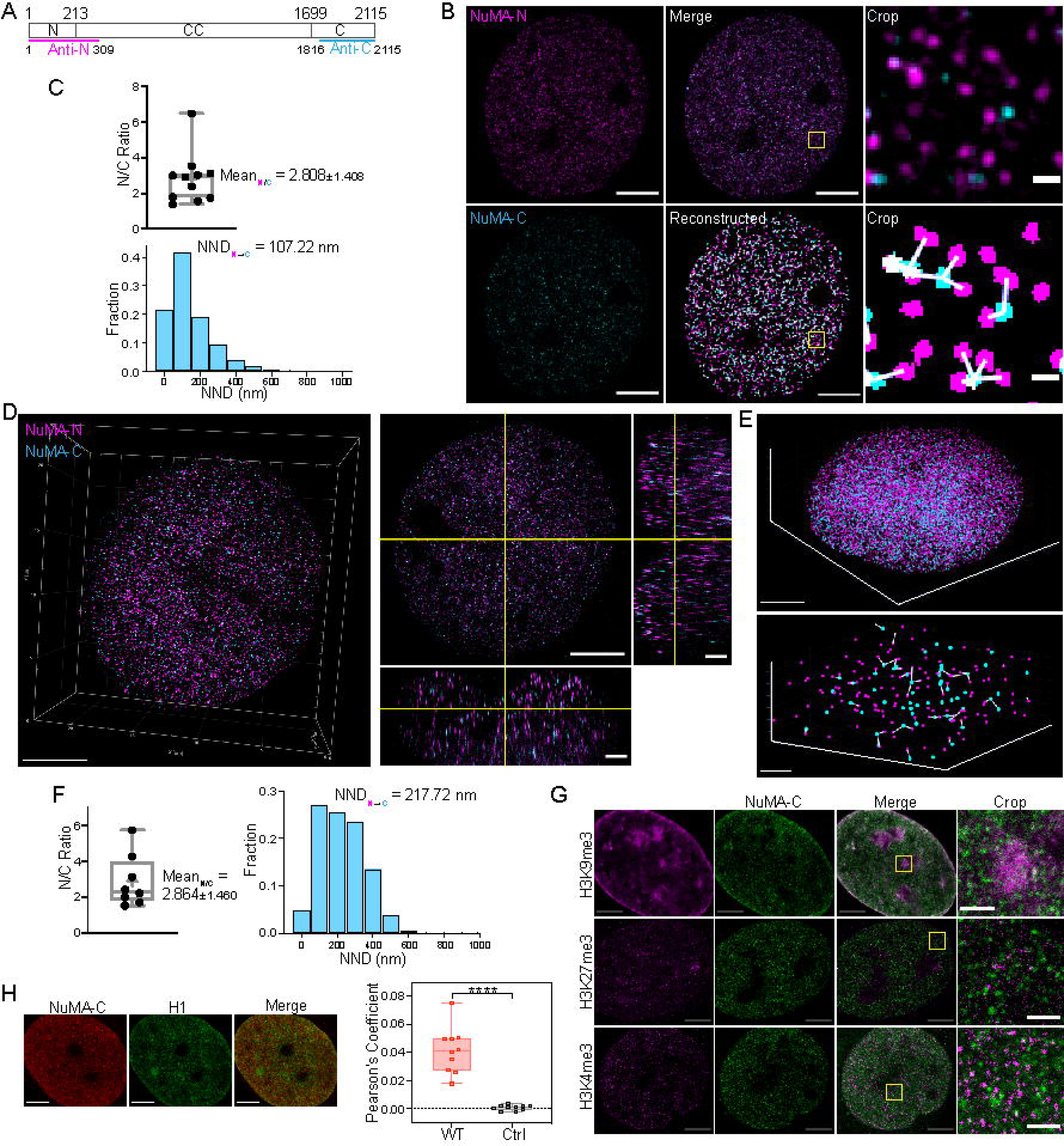
NuMA oligomerizes into quasi-network organization through its C-termini *in vivo*. A. Schematic diagram of antibody epitopes targeting NuMA-N and NuMA-C. B. Expanded 2D IF images of NuMA-N and NuMA-C and reconstructed NuMA oligomers in U2OS cells. NuMA-N was labeled by Atto647N (magenta) and NuMA-C was labeled by Alexa594 (cyan). Scale bars, 5 μm and 250 nm. C. Calculated ratio of numbers (upper) and distribution of nearest neighbor distance (NND) from NuMA-C to NuMA-N clusters in U2OS 2D expanded IF images (lower). D. Perspective view (left) and orthogonal view (right) of expanded 3D IF images of NuMA-N and NuMA-C in U2OS cells. Scale bars, 5 μm and 2 μm. E. Perspective 3D view and crop of reconstructed NuMA oligomers in U2OS 3D expanded IF images. Scale bars, 5 μm and 1 μm. F. Calculated ratio of numbers (left) and distribution of NND from NuMA-C to NuMA-N clusters in U2OS 3D expanded IF images (right). G. Expanded IF images of H3K9me3, H3K27me3 and H3K4me3 co-stained with NuMA-C in U2OS cells. Scale bars, 5 μm and 1 μm. Error bars represent SD (n=10). ****p < 0.0005, Mann-Whitney test. H. Expanded IF images of H1 co-stained with NuMA-C in U2OS cells (left) and Pearson’s coefficient of NuMA-C and H1 (right). The NuMA-C clusters were randomized and calculated as control. Scale bar, 5 μm.

The 2D characteristics of NuMA distribution were further validated in the three-dimensional space by 3D super-resolution imaging (Figure 5D-F). Notably, the mean NND from NuMA-C to NuMA-N in 3D images was 217.72 nm in U2OS cells (Figure 5F), which is close to the length of NuMA molecule and indicates that NuMA adopts an extended conformation in the nucleus. To prevent the influence of different dyes on efficiency of fluorescent labeling and imaging, we swapped fluorescent dyes for NuMA-N and NuMA-C, which yielded similar distribution patterns, stoichiometry and NND values (Figure S5B & S5D). We also performed the experiments and analysis in HCT116 cells and the results were similar with those obtained in U2OS cells (Figure S5C & S5E). Collectively, these imaging analyses supported that NuMA oligomerizes into quasi-meshwork organization through its C-termini *in vivo*.

### NuMA contributes to epigenetic maintenance of constitutive heterochromatin and repression of LTR expression

Since we have found that NuMA’s regulatory effect on chromatin architecture at the nucleosome level and facilitation effect on nucleosome stacking by promoting H1’s binding to DNA are mainly showed in regions enriched with heterochromatin markers, our subsequent investigations focused on alterations of heterochromatin markers induced by NuMA-depletion.

Because of the high abundance of NuMA in nucleus, we utilized expansion microscopy to characterize the localization of NuMA-C and heterochromatin markers, H3K9me3 and H3K27me3, which marks constitutive heterochromatin and facultative heterochromatin, respectively. H3K4me3, the transcriptionally active euchromatin marker, was also labeled as negative control. Among the three epigenetic markers, we only observed an obvious distribution pattern around H3K9me3 speckles for NuMA-C (Figure 5G). Next, as the expression level of all these markers remained stable (Figure S6B), the distribution of H3K9me3 became more dispersed in the nucleoplasm after NuMA-depletion in HCT116 cells, as rated by the heterogenous index (Described in “Immunofluorescence staining” in Methods). In contrast, the distribution of H3K27me3 remained unperturbed upon NuMA-depletion, which is similar to H3K4me3 (Figure 6A). These results imply that while the effect of NuMA-depletion on chromatin architecture and nucleosome stacking is most significant in heterochromatin, its influence on heterochromatin markers is specifically concentrated on H3K9me3, the constitutive heterochromatin modification. Similar results were obtained in NuMA-depleted U2OS cells (Figures S6A and S6C), reinforcing the specificity of NuMA’s role in regulation of constitutive heterochromatin modification.

**Figure 6.**
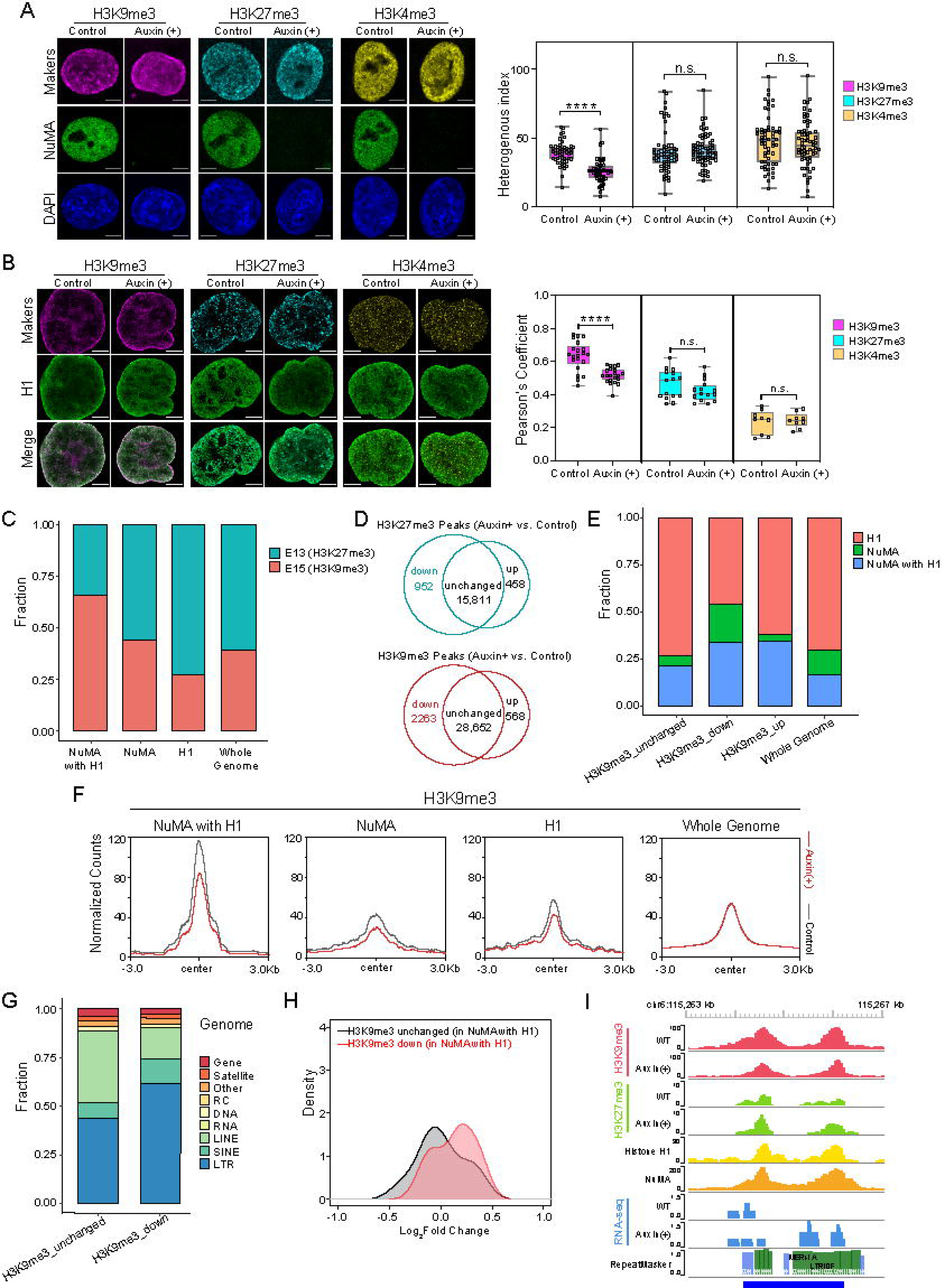
NuMA contributes to epigenetic maintenance of constitutive heterochromatin and repression of LTR expression. A. IF imaging (left) and quantification of the heterogenous index (right) of H3K9me3, H3K27me3 and H3K4me3 in HCT116-mAID-NuMA cells after induced by auxin, with untreated HCT116-mAID-NuMA cells as control. Error bars represent SD (n ≥50). ****p < 0.0005, Mann-Whitney test. Scale bar, 5 μm. B. Representative expanded IF images of histone H1, NuMA co-stained with H3K9me3, H3K27me3 and H3K4me3 (left) in NuMA-depleted HCT116-mAID-NuMA cells and untreated HCT116-mAID-NuMA cells as control. Pearson’s coefficient of histone H1 with H3K9me3, H3K27me3 and H3K4me3 (right). Scale bar, 5 μm. C. Fraction of H3K9me3 and H3K27me3 enrichment in genomic regions where NuMA-C and H1 bind and overlap, and among the whole genome. D. Percentage of genomic regions with up-, down- and unchanged peaks of H3K27me3 (upper) and H3K9me3 (lower) upon NuMA-depletion. E. Fraction of NuMA-C and H1 enrichment in genomic regions with unchanged H3K9me3, down-regulated H3K9me3, up-regulated H3K9me3 upon NuMA-depletion and in the whole genome. The peaks were aligned using the center of the peaks. F. Changes of H3K9me3 CUT&Tag peaks upon NuMA-depletion in different categories (regions where NuMA-C and H1 are co-bound and regions bound by NuMA or H1 alone), with the change profile in the whole genome as control. G. Percentage of multiple types of transposable elements in all genomic regions with unchanged H3K9me3 and down-regulated H3K9me3 upon NuMA-depletion. H. Expression changes of LTRs in NuMA-C and H1 co-bound regions with unchanged and down-regulated H3K9me3 upon NuMA-depletion. I. Representative example of expression up-regulation in genomic regions where NuMA-C and H1 are co-bound and with down-regulated H3K9me3 upon NuMA-depletion.

Considering that NuMA maintains nucleosome stacking through its interaction with H1 (Figure 4) and H1 is widely known to be correlated with heterochromatin formation and maintenance^43–46^, we assessed the spatial distribution of H1 alongside NuMA-C and the epigenetic markers. We observed a strong spatial correlation between NuMA-C and H1 (Figure 5H), consistent with their co-localization on the chromatin (Figure 3D and 3E). Moreover, NuMA-depletion was found to greatly reduce the co-localization degree between H1 and H3K9me3, but had no effects on H3K27me3 or H3K4me3 (Figure 6B), further substantiating NuMA’s specific regulatory effect on constitutive heterochromatin modification.

After observing the spatial distribution of heterochromatin markers, we applied CUT&Tag to profile the heterochromatin markers, H3K9me3 and H3K27me3, along the whole genome. In untreated HCT116 cells, H3K9me3 accounted for a higher proportion at binding sites that NuMA-C and H1 overlapped than H3K27me3 (Figure 6C), which suggested that NuMA’s effect on H1 is more concentrated in constitutive heterochromatin than facultative heterochromatin. Then we measured changes of the enrichment of H3K9me3 and H3K27me3 on chromatin after NuMA-depletion, and found that the number of down-regulated H3K9me3 peaks was the highest among all of the altered repressive markers (Figure 6D), illustrating that NuMA-depletion mainly causes down-regulation of constitutive heterochromatin modification. Reciprocally, the overlap of NuMA-C and H1 also located more in regions with down-regulated H3K9me3 than regions with up-regulated and unchanged H3K9me3 (Figure 6E and S6D). When summarizing H3K9me3 peaks according to the binding sites of NuMA-C and H1, we found that the down-regulation of H3K9me3 was most significant at regions where NuMA-C and H1 overlapped (Figure 6F), which suggests that NuMA coordinates with H1 in maintenance of constitutive heterochromatin modification. On the contrary, the absence of apparent pattern in the enrichment of H3K27me3 (Figure S6E) and summary of H3K27me3 peaks in binding sites of NuMA overlapping H1 (Figure S6F) also support this conclusion.

Previous research reported that linker histone H1 is enriched in the constitutive heterochromatin that silence repetitive elements in mESCs, and that acute deletion of H1 leads to substantial derepression of repetitive element gene expression^45^. Then we were intrigued by NuMA’s potential effects on transcription, especially in regions that NuMA and H1 overlap. RNA-seq analysis revealed that NuMA-depletion minimally affected gene transcription across the whole genome (Figure S6G) with no specific Gene Ontology (GO) enrichment (Figure S6H), which are consistent with that NuMA mainly regulate constitutive heterochromatin with relative low gene content. However, when we turned to non-coding genomic regions, we found that upon NuMA-depletion, the long terminal repeat (LTR) elements accounted for higher proportion in regions with down-regulated H3K9me3 than regions with unchanged H3K9me3 (Figure 6G). Based on this, the transcription level of LTRs in regions that NuMA and H1 overlapped were also altered, which was significantly increased with up-regulated H3K9me3 (Figure 6H, 6I). These results indicated that NuMA collaborates with H1 to contribute to epigenetic maintenance of constitutive heterochromatin and repression of LTR expression, strongly suggesting NuMA’s crucial role in cell fate regulation during the interphase, such as cell differentiation and senescence.

## Discussion

Based on the data presented, we proposed a hypothetical model that NuMA serves as nucleoskeleton protein to regulate heterochromatin maintenance and genome organization in the interphase. NuMA interacts with linker histone H1 and stabilizes H1’s enrichment on chromatin to facilitate nucleosome stacking. Besides of its interplay with H1, NuMA also oligomerizes into quasi-meshwork organization through its C-termini. Put together, both NuMA-H1 and NuMA-NuMA interactions are coordinated in promoting constitutive heterochromatin compaction and repression of non-coding LTRs (Figure 7).

**Figure 7.**
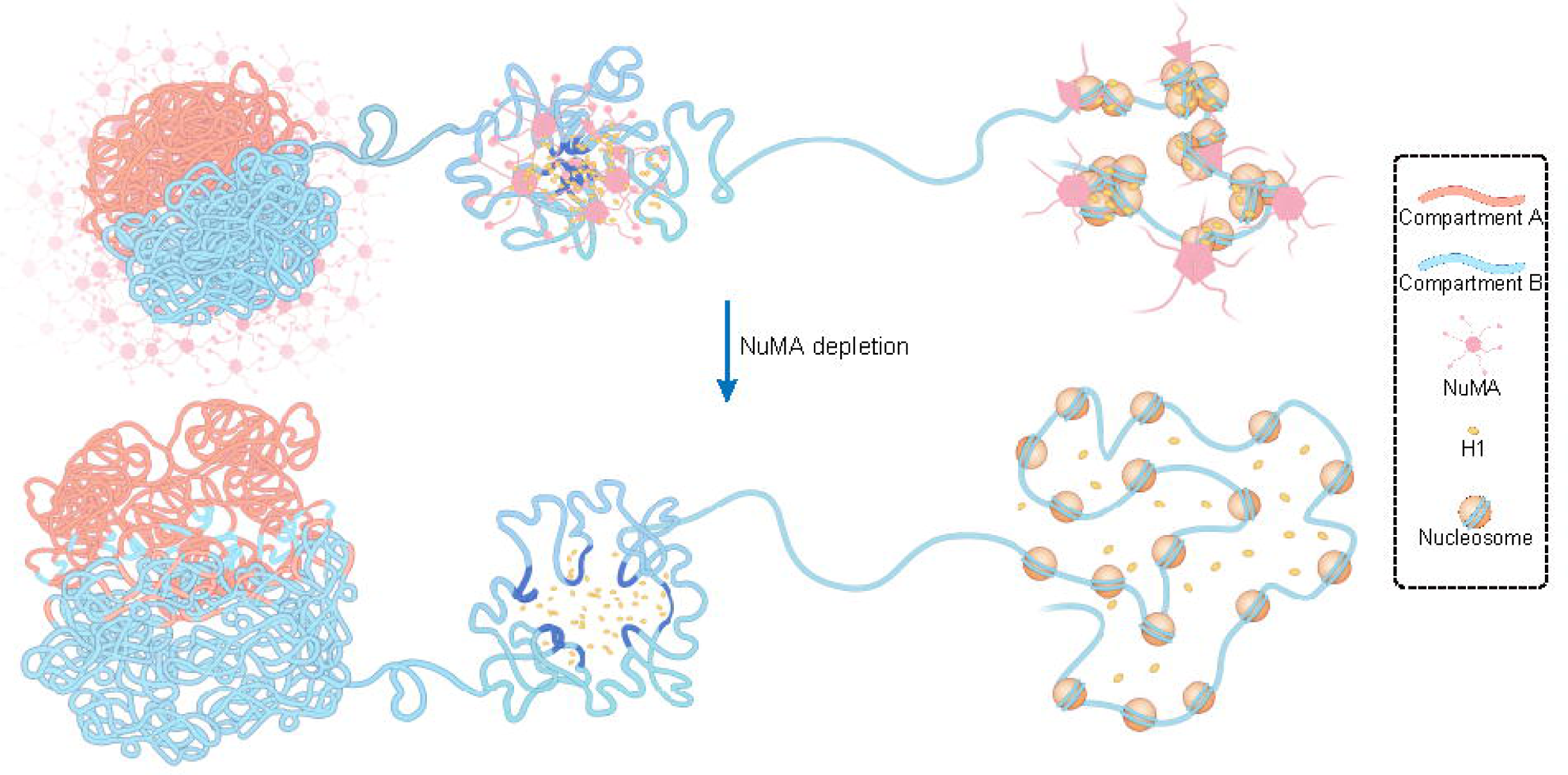
Models for NuMA’s promotion on constitutive heterochromatin compaction through stabilizing linker histone H1. NuMA interacts with H1 with its C-terminus, stabilizes H1’s binding to DNA, enhance H1’s chromatin compaction effect and nucleosome stacking, then contributes to epigenetic maintenance of constitutive heterochromatin and repression of LTR expression.

Beyond the conventional classification of euchromatin and heterochromatin, the 3D hierarchical organization of chromatin introduces an additional regulatory dimension governing fundamental biological processes, such as DNA replication, transcription, and cell fate decision. Meanwhile, chromatin’s hierarchical architecture exhibits profound spatiotemporal coupling with euchromatin and heterochromatin. Genomic regions of Compartment A are enriched with euchromatin epigenetic markers and transcriptionally active, while Compartment B is mostly transcriptionally repressive and enriched with heterochromatin markers^47^. Combining Hi-C and ATAC-seq, we found that NuMA-depletion mainly causes B-to-A compartment transition (Figure 1F-I) and increased genomic accessibility of regions enriched with repressive epigenetic markers (Figure 2A-B, S2A-D), demonstrating that NuMA regulates heterochromatin compaction. These insights also significantly enhance our understanding of the nucleoskeleton’s role in shaping chromatin architecture in mammalian cells.

Besides of the compartment-level reorganization, the nucleosome level of chromatin also exhibits significant alterations induced by NuMA-depletion, which is reflected by shortened NRL and dispersed nucleosome clutches (Figure 2, S2). Given that linker histone H1 is one of the direct regulatory factors of NRL^27, 37^, we confirmed the specific interaction between NuMA and H1, and proved that NuMA stabilizes H1’s enrichment on chromatin to facilitate nucleosome stacking (Figure 3, 4 and S3, 4). This finding correlates nucleoskeleton and chromatin architecture at the level of nucleosomes for the first time. Intriguingly, the phenotypes induced by NuMA-depletion, including B-to-A compartment transition, chromatin decompaction, shortening of NRL, dispersed nucleosome clutches and dispersion of H3K9me3 in heterochromatin regions, are similar with those induced by H1 TKO reported previously^33, 34, 44, 45^. This observation is also in line with the enrichment of H1 in heterochromatin regions^33^. Moreover, in contrast to the changes in nucleosome stacking, chromatin structure at the TAD level, manifested by the distribution of TAD length and insulation scores around TAD boundaries, remained unchanged in control and NuMA-depleted cells (Figure S1H and S1I). Along with the highest degree of colocalization between NuMA and H1 in heterochromatin region (Figure 3F) and the lack of conserved sequence features at NuMA binding sites (data not shown), these results all implies NuMA’s reliance on H1 for chromatin binding and 3D genome organization.

Moreover, *in situ* cryo-ET results provided direct structural evidence that NuMA stabilizes the binding of histone H1 to DNA and nucleosomes (Figure 4 and S4). The increased NND among nucleosomes observed in cells overexpressing NuMA-dC compared to those expressing NuMA-FL suggests weaker nucleosome stacking and less compact chromatin organization, which are consistent with the phenotypes induced by NuMA-depletion (Figure 1, 2, S1, and S2). In addition, the absence of H1 density in NCPs map from cells overexpressing NuMA-dC further supports the essential role of NuMA-C in stabilizing H1’s association with nucleosomes. Previous researches reported that H1 condensates with DNA and nucleosome fibers to promote heterochromatin formation^43, 45, 46, 48^. Besides, H1 also functions in heterochromatin formation and maintenance through interaction with other proteins which in turn modify chromatin or take part in DNA-based processes, like recruiting histone methyltransferase Su(var)3-9 to chromatin and providing binding platform for HP1 in Drosophila melanogaster^43^. Therefore, the regulatory effects of NuMA on heterochromatin are predominantly attributable to its stabilization of histone H1, thereby unveiling a novel layer of mechanisms by which nucleoskeleton proteins regulate heterochromatin formation and maintenance.

The formation and maintenance of heterochromatin are critical for early embryonic development and differentiation processes, during which programmed gene silencing ensures precise spatiotemporal control of developmental cues^1–5^. Besides of nucleosome spacing tuning mentioned above, H1 also participates in programmed gene silencing in development. For example, when the protein level of the single somatic H1 subtype was reduced to around 20% of the normal content by RNAi, the H1-depleted D. melanogaster larvae wouldn’t develop to adult flies^49^. Besides, H1 was also reported to contribute to repression of TEs. In D. melanogaster, loss of H1 caused derepression of more than 50% of TEs and only about 10% of protein-coding genes by physically tethering Su(var)3-9 to facilitate methylation of H3K9^43^. This function of H1 to contribute to TE silencing by participating in constitutive heterochromatin formation well matches down-regulation of H3K9me3 peaks induced by NuMA-depletion and the higher co-localization proportion of H3K9me3 with overlapped NuMA-C and H1 on genome than H3K27me3. LTR belongs to the retrotransposons of TE family, and is implicated in several critical biological processes, such as generating mutations in evolution. Typically, most TEs are inactive and tightly regulated to ensure genomic stability^50, 51^. In 2022, Gao and colleagues reported that LTRs are gradually silenced by stage-specific H3K9me3 establishment in human early embryos^52^. In 2023, Liu and colleagues found that derepression of endogenous retrovirus (ERVs), which accounts for ∼97% of LTR, is associated with epigenetic derepression of H3K9me3 and contributes to cellular senescence^53^. Since the establishment and maintenance of H3K9me3 are critical for repression of LTRs, NuMA’s compaction effect on constitutive heterochromatin indicates its nonnegligible roles in the maintenance of genomic stability and cell fate decision. Through interrogation of public database^54–56^, we found that NuMA’s expression level is stage-specifically elevated during embryogenesis (data not shown), providing orthogonal validation of our proposition that NuMA plays important roles in cell fate decision, rather than merely functioning as a mitotic regulator.

Dispersed nucleosome clutches induced by NuMA-depletion also indicates a potential role for NuMA in cellular differentiation and development. The concept of nucleosome clutch was first put forward in 2015^34^. Using STORM imaging, the authors found that nucleosomes are arranged in heterogeneous groups of around 20-100 nm diameter along the chromatin fiber. Importantly, a correlation was discovered between the spatial distribution, size, compaction of nucleosome clutches and cell pluripotency. For example, ground-state stem cells have lower-density clutches containing on average only a few nucleosomes than differentiated ones. Therefore, the dispersion and decrease of nucleosome clutches induced by NuMA-depletion (Figure 2D & S2H) and increase of NND among nucleosomes induced by overexpressing of NuMA-dC (Figure 4E) indicate that NuMA has the potential in regulation of differentiation and development processes.

In summary, we have provided comprehensive evidence proving that NuMA promotes constitutive heterochromatin compaction and consequently represses LTR expression by stabilizing H1 on chromatin in the interphase. Besides, the NuMA meshwork is probably another mechanism by which it’s involved in chromatin organization, like imposing spatial constraints on chromatin topology.

There are some intriguing questions awaiting to be answered. For instance, a few recent studies have demonstrated that H1 can condensate with DNA *in vitro* through its highly disordered C-terminal tail^57^, and H1 promotes chromatin’s liquid-liquid phase separation (LLPS) *in vitro*^46, 48, 58^. Besides, NuMA-C was also reported to regulate mitotic spindle assembly and structural dynamics *via* phase separation^59, 60^. Thus, the mechanism of NuMA’s stabilization effect on H1 in the interphase is also probably related to phase separation. On the other hand, as an essential mitotic spindle regulator, it is unclear how the different functions of NuMA from the mitotic phase to the interphase are transitioned. One possibility is that post-translational modifications of NuMA control the function transition, like phosphorylation. As a hint, NuMA binds both microtubules and H1 through its C-terminus, and the most important phosphorylation modification of NuMA after entering mitotic phase is also exactly located in H1BD2^19^. Besides 3D genome organization, NuMA probably also contributes to nuclear mechanotransduction because of its unique conformation and oligomerization mode, which would be tempting for further exploration.

## Methods

### Cloning

The sgRNA sequence (5’-GGTGGCGTGGAGTGTCATCT-3’) targeting to *NuMA*’s N-terminal locus was generated using the online CRISPR design tool (http://crispor.tefor.net). The sgRNA sequence (5’-GGTGGGGCCACTCACTGGTA-3’) targeting to *NuMA*’s C-terminal locus was referred to previous report^61^. The sgRNA fragments were annealed from commercially synthesized single-strand oligos and then inserted into the pX330-U6-Chimeric_BB-CBh-hSpCas9 vector (Addgene, 42230) using Golden Gate cloning (BbsI-HF, NEB, R3539M).

The DNA fragment of NuMA and histone H1 isoforms was amplified from a cDNA library produced from HCT116 cells by reverse transcription. Truncations of NuMA was generated through PCR amplification and then cloned into pEGFP-N1, pET-28a (+) and lentivirus vectors through Gibson assembly.

### Cell culture and transfections

The human colon tumor cell line HCT116 was cultured in McCoy’s 5A medium (Gibco, 16600082), while HEK293 cells and human cancer cell lines U2OS and HeLa were cultured in DMEM medium (Gibco, C11995500BT). All types of medium contain 10% fetal bovine serum (Gibco, 10091148) and 1% penicillin-streptomycin (Gibco, 15140122), and all cells were cultured in a humidified incubator at 37 °C with 5% CO_2_. These cells were tested and found to be free of mycoplasma contamination. Before rapid degradation of NuMA, cells were synchronized to the G1/S boundary by two rounds of blocking as described previously^62, 63^. In the first round of synchronization, cells were treated with 2 mM thymidine (Sigma-Aldrich, T1895) for 15 hours followed by releasement to fresh medium supplemented with doxycycline (1 μg/ml for HCT116 cells and 5 μg/ml for U2OS cells, Beyotime, ST039) for 10 hours. Then in the second round of synchronization, cells were treated by 2 μg/mL aphidicolin (Sigma-Aldrich, A4487) for 15 hours. NuMA-depleted cells were treated with 500 μM auxin/IAA (Sigma-Aldrich, I2886) for 7 hours at the end of the second blocking period.

For transient transfection, cells were passaged to approximately 60% confluency one day ahead. Transfection was performed using Neofect transfection reagent (Neofect, TF20121201) following the manufacturer’s protocol. Approximately 0.5 μg plasmids were transfected into cells in 12-well plates, 1 μg plasmids were transfected into cells in 35 mm Petri dishes or 6-well plates, 2 μg plasmids were transfected into cells in 60 mm dishes and 10 μg plasmids were transfected into cells in 100 mm dishes. After transfection, cells were cultured for 72 hours until the samples were imaged or co-IP experiments.

For over-expression of NuMA and its truncations, transfection was accomplished by lentiviral infection. Lentivirus was prepared referring to officially recommended protocols of Addgene (https://www.addgene.org/protocols/lentivirus-production/). HEK293T cells were passaged to approximately 60% confluency one day ahead in 100 mm plates. 10 μg lentiviral transfer plasmids, 5 μg lentiviral packaging plasmid psPAX2 (Addgene, 12260) and 7.5 μg envelope expressing plasmid pMD2.G (Addgene, 12259) were co-transfected. The lentiviral particles were harvested by filtering the supernatant through a 0.45 μm filter 24, 48 and 72 hours post transfection and either used immediately or stored at -80 °C.

### Knock-in by CRISPR/CAS9

Construction of HCT116 and U2OS cells expressing OsTIR1 was performed as reported previously^28^. Plasmids AAVS1 T2 CRIPR in pX330 (Addgene, 72833) and pMK243 (Tet-OsTIR1-PURO) (Addgene, 72835) were co-transfected and positive clones were screened by puromycin. Then pX330-based CRISPR/CAS plasmid which contains sgRNA targeting to *NuMA’*s N-terminus and homology-directed repair template plasmid containing mAID and Clover were co-transfected into the Tet-OsTIR1 cells. After around two days of culture, cells were sorted *via* FACS to isolate single cells expressing the Clover protein for imaging. After two weeks, single-colony expansion was verified using western blots. Similar procedures were performed to generate HCT116 cells with N-terminal Avitag and eGFP and C-terminal HA tag in the *NuMA* gene locus respectively.

### Antibodies

The antibodies used are listed in Supplementary Table 1.

### Western-blotting

For whole-cell samples, cells were harvested by centrifugation at 200 g for 3 min and lysed in RIPA buffer supplemented with 0.1 mg/mL PMSF for 10 minutes on ice. Cell lysate, immunoprecipitation and pull-down input and product were added with SDS-page loading buffer and boiled at 98 °C for 10 minutes. Then samples were run on 10% or 15% polyacrylamide gel and wet-transferred to 0.45 μm PVDF membrane. Next, the membrane was blocked with 5% skim milk dissolved in TBST buffer (Tris-HCl 10 mM, pH 7.5, NaCl 150 mM, Tween-20 0.05%) for 1 hour at room temperature, incubated with primary antibodies at 4 °C overnight. After washed three times with TBST for 5 minutes at RT with shaking, the membrane was incubated with HRP-conjugated secondary antibodies for 1 hour at RT. Finally, the membrane was washed with TBST for five times and covered with ECL substrate. The blot images were acquired with Amersham Imager 600 (GE).

### Immunofluorescence staining

Cells were fixed with 4% paraformaldehyde in 1× PBS for 10 minutes and permeated with 0.5 % Triton-100 in 1× PBS for 30 minutes at RT. Then samples were blocked with blocking buffer (1× PBS, BSA 5%, Triton-100 0.5 %) for 1 hour at RT, and incubated with primary antibodies diluted in blocking buffer at 4 L overnight. The next day, samples were washed five times with 1× PBS for 5 minutes at RT with shaking and incubated with dye-conjugated secondary antibodies diluted in blocking buffer at RT for 1 hour. After washed five times, samples were post-fixed with incubated with 4% paraformaldehyde in 1× PBS for 10 minutes at RT. Finally, samples were stained with 1 μg/mL Hoechst 33342 diluted in 1× PBS for 10 minutes at RT. Cells in 35 mm Petri dishes were kept away from light in 1× PBS at 4 L until imaging. Cells on coverslips were mounted with Fluoromount-G and dried in the shade, and kept away from light at 4 L until imaging. Images were acquired using TCS SP8 STED 3X (Leica) microscope with a ×100 objective and post-processed using Fiji (ImageJ) and MATLAB.

The heterogeneous index was calculated to reflect the uniformity of distribution like histone modifications. Before batch processing, nucleoli and extranuclear regions were manually selected and the intensity of these parts were set to 0. Then average intensity of each image was set to same value to eliminate variations caused by sample preparation and imaging. Considering the shape of the aggregate structures which form around 5-pixel thick sticks, we used a 5×5-pixel box traversing the image and calculated the sum of intensity inside the box. The values of 0 were left out, and the histogram of frequency distribution was calculated and fitted with double Gaussian distribution. The two Gaussian curves represented the distribution of sparse regions and dense regions respectively. Distance between the centers of two Gaussian peaks was calculated to quantify the aggregation level. It should be noted that the rationality of fitting results is closely related to the starting point of fitting. Therefore, the position with the highest frequency was used as the default starting point of the double Gaussian fitting, and the starting point was adjusted manually for the results obviously not reasonable, such as negative Gaussian curve strength.

### Fluorescence *in situ* hybridization (FISH)

Cells used for FISH labeling were cultured on coverslips in 12-well plates. The whole experiment procedure was according to Metasystems’ protocols. After washed with pre-warmed 1× PBS, cells were fixed in pre-cooled 3:1 methanol/acetic-acid for 10 minutes at -20 L. The fix reagent was removed and the coverslip was air-dried. FISH probe targeting to chromosome 2 and 18 (Metasystems, D-0302-100-FI XCP 2 and D-0318-100-OR XCP 18) were mixed with 5 μL each and dropped on a glass slide. The coverslip with cells was inverted onto the probes and sealed with nail polish. Then the sample and probes were heated at 75 L for 2 minutes to be denatured. Next, the sample was incubated in a humidified chamber at 37 °C overnight. The next day, the coverslip was removed and washed in 0.4× SSC buffer (pH 7.0) at 72 °C for 2 minutes. Then the sample was washed in 2× SSC buffer supplemented with 0.05% Tween-20 at RT for 30 seconds. Finally, the sample was stained with 1 μg/mL Hoechst 33342 diluted in 1× PBS for 10 minutes at RT, mounted with Fluoromount-G, dried in the shade, and kept away from light at 4 L until imaging. Images were acquired using TCS SP8 STED 3X (Leica) microscope with a ×100 objective and post-processed using Fiji (ImageJ).

### DNase I /Mnase digestion assays

Cells (approximately 2 ×L10^6^) were collected by centrifugation at 200 *g* for 3 minutes. After washed by cold 1× PBS twice, cells were suspended and lysed by nucleus extraction buffer (Tris-HCl 10 mM, pH 7.5, NaCl 10 mM, MgCl_2_ 3 mM, IGEPAL CA-630 0.1% and 0.1 mg/mL PMSF) on ice for 60 seconds. Then centrifuge immediately at 500 *g* for 10 minutes at 4 L to pellet nuclei and remove supernatant completely. Extracted nuclei were resuspended with 600 μL DNase I reaction buffer (Tris-HCl 10 mM, pH 7.5, MgCl_2_ 2.5 mM, CaCl_2_ 0.1 mM, sucrose 0.3M) or Mnase reaction buffer (Tris-HCl 10 mM, pH 7.5, NaCl 10 mM, KCl 10 mM, MgCl_2_ 3 mM, CaCl_2_ 10 mM, sucrose 0.3M) and divided into six tubes. Each tube was treated with different units of DNaseI (Thermo Fisher, EN0523) or Mnase (Thermo Fisher, EN0181) at 37 L and 800 rpm rotation for 10 minutes. The reactions were terminated by 50 μL 0.5 mM EDTA (pH 8.0) and 50 μL 10% SDS. Then, equal volume of 2× DNA extraction buffer (Tris-HCl 20 mM, pH 8.0, NaCl 200 mM, EDTA 100 mM, SDS 2%) was added to each tube and samples were digested with 20 μg/mL RnaseA (TransGene, GE101) at 37 L for 30 minutes and 0.2 mg/mL Proteinase K at 65 L for 1 hour. Finally, DNA was extracted with equal volume of phenol/chloroform. Equal amounts of DNA (200 ng) were resolved on 1% agarose gel and stained with EB. High MW chromatin was measured by ImageJ.

### In situ Hi-C

The *in situ* Hi-C libraries were prepared and sequenced as the published method^30^. Briefly, cells were grown to approximately 70–80% confluency, washed with 1×PBS, crosslinked using 1% formaldehyde, and suspended in Hi-C lysis buffer (Tris-HCl 10 mM, pH8.0, NaCl 10mM, IGEPAL CA-630 0.2%) supplemented with protease inhibitors. 100 Units DpnII restriction enzyme was added *per* sample at 37 L with rotation for overnight chromatin digestion. DNA ends were marked with biotin and then ligated together *in situ*. After crosslink reversal, the DNA was sheared to 300∼500 bp fragments, and then biotinylated ligation junctions were recovered with streptavidin beads. Hi-C libraries were amplified using PCR. After that, sequencing adaptor was added. All sequence data were produced using Illumina HiSeq X Ten paired-end sequencing.

Raw reads were firstly cut adaptor and filtered to generate cleaned fastq files using TrimGalore (https://www.bioinformatics.babraham.ac.uk/projects/trim_galore/). Next, Hi-C data preprocessing was done using HiC-Pro^64^. Briefly, reads were first aligned on the hg19 reference genome. Uniquely mapped reads were normalized using Iterative Correction and Eigenvector (ICE) decomposition and library size.

For compartment A/B analysis, HiTC^65^ was used to calculate the PC1 (at 150-kb resolution) with custom R script. The whole genome was classified into two compartments, A and B compartment, based on the positive or negative PC1 values. The compartment with higher gene density while the B compartment with lower gene density. To investigated compartment switching, we compared the PC1 values between control and NuMA depleted HCT116 cells, using zero as the PC1 cutoff. We defined the delta PC1 to describe A-to-B or B-to-A compartment switching, compaction and decompaction:

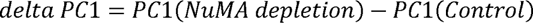

If one compartment had the delta PC1 ≥ 0.005 and the sign of PC1 (NuMA-depletion) and PC1 (Control) were opposite, it would be defined as B-to-A compartment switching; if one compartment had the delta PC1 ≤ -0.005 and the sign of PC1 (NuMA-depletion) and PC1 (Control) were opposite, it would be defined as A-to-B compartment switching; if one compartment had the delta PC1 ≥ 0.005 and the sign of PC1 (NuMA-depletion) and PC1 (Control) were identical, it would be defined as compartment decompaction; if one compartment had the delta PC1 ≤ -0.005 and the sign of PC1 (NuMA-depletion) and PC1 (Control) were identical, it would be defined as compartment compaction. Juicebox (http://aidenlab.org/juicebox/) was used to visualize Hi-C contact matrix^66^. IGV (https://igv.org) was used to display compartment switching.

### ATAC-seq

Each sample (approximately 5L×L10^5^ cells) was centrifuged with 500 g, 4 °C for 5 minutes to collect cell pellet. The cell pellet was resuspended with moderately cold lysis buffer, and library construction used the TruePrep DNA Library Prep Kit V2 for Illumina (Vazyme, TD501). DNA libraries were produced after 12 cycles of PCR amplification using the TruePrep library prep kit. After purification, the paired-end sequencing was performed on the Illumina Novaseq 6000 platform.

Sequencing adaptors were removed from the raw ATAC-seq reads using TrimGalore (https://www.bioinformatics.babraham.ac.uk/projects/trim_galore/), and the clean data were mapped to the human reference genome (hg19) using Bowtie2 (v.2.4.1)^67^. Picard (https://broadinstitute.github.io/picard/) was used to remove PCR duplicates.

The nuclear repeat length (NRL) was calculated using NRLfinder as previous publication^33^. Briefly, read lengths were extracted and converted into a frequency histogram, which was then smoothed using a digital 6th-order Butterworth filter with a zero-phase shift and a cutoff frequency of 0.04 cycles/read. This cutoff was empirically optimized to reduce noise from mononucleosomal DNA winding artifacts. Local minima and maxima were identified from the first derivative of the filtered histogram, with the second peak maximum corresponding to the dinucleosomal periodicity. The NRL shift between conditions (e.g., control vs. NuMA-depleted HCT116 cells) was calculated the mean difference between the first two peak maxima of each sample. All analyses were performed in Python 3.9 with NumPy, SciPy, and Matplotlib libraries.

For chromatin-state modelling, we used the ChromHMM (v.1.19)^32^. The input data of ATAC-seq and RNA-seq reported in this manuscript was generated as described above. Additional input data including ChIP-seq for CTCF, H3K4me3, H3K27me3, H3K4me1, H3K36me3 and H3K9me3 were download from ENCODE (https://www.encodeproject.org). briefly, raw bam files were download and replicates were combined. BinarizeBam and LearnModel tools in ChromHMM was used to generate chromatin state model with default settings. Emissions parameters were visualized in R.

For LAD annotation of ATAC-seq peaks, the public data of LaminB1 DamID of HCT116 was download from 4D Nucleome Data Portal (https://data.4dnucleome.org/) and analyzed as previous publication^9^.

### CUT&Tag

The CUT&Tag assay was performed using the NovoNGS® CUT&Tag 3.0 High-Sensitivity Kit (NovoProtein, N259-YH01). 1L×L10^5^ cells were washed twice with 1.5 mL of wash buffer and then mixed with activated concanavalin A beads. Primary antibody targeting histone H1 was added to the system for 2 h at room temperature. After successive incubations with the primary antibody and secondary antibody at RT for 1.5 hours. The cells were washed and incubated with pAG-Tn5 for 1.5Lh. Then, tagmentation buffer was added to activate tagmentation for 1 h. The tagment DNA fragments were purified using tagment DNA extract beads and washed by 80% ethanol. DNA libraries were produced after PCR amplification and sequenced using the Illumina NovaSeq 6000 platform.

TrimGalore (https://www.bioinformatics.babraham.ac.uk/projects/trim_galore/) was used to cut adaptors, and trimmed reads were aligned to the human genome (hg19) using Bowtie2 as described in ATAC-seq data analysis. Reads were sorted and converted to BAM format and duplicates were marked using Picard (https://broadinstitute.github.io/picard/) using the *MarkDuplicates* module. After normalizing the samples to the same sequencing depth, deepTools2 (v 3.3.1) software^68^ was used to plot the heat maps to show signals around peak regions with default parameters. Peaks were called using Macs2 with a relaxed *q* value threshold of 0.001.

For peak calling, MACS2^69^ (v2.2.7.1) was employed to identify enriched regions with the *q* value threshold of 0.001. Blacklisted regions (from the ENCODE Project) were excluded using bedtools^70^ (v2.30.0). Peak regions were then normalized to the same sequencing depth using deepTools2^68^ (v3.3.1), and heat maps of signal intensity around peak summits were generated with the *computeMatrix* and *plotHeatmap* functions, setting the region of interest to ±3 kb from peak centers.

To analyze co-localization with histone modifications, peak intersections were performed using bedtools *intersect* with a minimum overlap of 1 bp. Genomic annotation of peaks was conducted using HOMER (v4.11.1). For visualization, IGV^71^ was used to generate genome browser tracks, and statistical analyses were performed in R with custom scripts.

### RNA-seq

Total RNA was extracted using the MolPure Cell RNA Kit (YEASEN, 19231ES50). RNA sequencing libraries were constructed using the NEBNext Ultra RNA Library Prep Kit for Illumina (NEB, E7530). RNA-seq paired-end reads were sequenced on the Illumina NovaSeq 6000 platform. The raw RNA sequences were cleaned using TrimGalore (https://www.bioinformatics.babraham.ac.uk/projects/trim_galore/) and mapped to human reference genome hg19 by STAR (v2.7.1a) with default parameters. All mapped bam files were converted to bigwig using bedtools (v2.24.0)^70^ for visualization in IGV (https://igv.org). High-quality mapped reads were quantified using htseq-count (v0.11.3)^72^. Differentially expressed genes were analyzed by DEseq2^73^. GO enrichment analysis was performed using Enrichr^74^. The integrated analysis of RNA-seq data and Hi-C data was done in custom R scripts.

### Super-resolution imaging and live-cell imaging

STED imaging was performed using a commercial STED microscope (TCS SP8 STED 3X, Leica Microsystems, Germany) equipped with an HCX PL APO ×100/1.40 NA objective. Alexa 594 labeled samples were excited with white laser at the wavelength of 594 nm, and Atto 674N labeled samples were excited with white laser at the wavelength of 647 nm. STED laser for Alexa 594 was continuous wave at the wavelength of 660 nm, and the wavelength of STED laser for Atto 674N was 775 nm pulsed laser. All images were acquired with the LAS AF software (Leica). De-convolution was performed with LAS AF software (Leica) and post-processed using Fiji (ImageJ).

HeLa cells co-transfected with the eGFP-H1.1 and different NuMA truncations fused with mScarlet were passaged into 35 mm Petri dishes to 60-70 % confluency. Imaging was performed using a spinning disk confocal system (Live SR CSU W Nikon) with an EMCCD (iXon DU-897E) mounted on a Nikon Ti-E microscope with a CFI Apo TIRF ×100 Oil (N.A. 1.49) objective. During image acquisition, the cells were incubated at 37 °C with 5% CO_2_. All images were post-processed using Fiji (ImageJ).

### Expansion sample preparation and image acquisition

To prepare hydrogel samples, we performed the gelation processes referring to Joerg Bewersdorf and the co-workers’ instructions^41, 42^. Before gelation, fixed cells were labeled with first antibody and second antibody as described above. Then the samples were fixed with EM-grade 3% PFA (Electron Microscopy Sciences, 157-8) and 0.1% glutaraldehyde (GA, Electron Microscopy Sciences, 16020) in 1× PBS for 15 min at RT with rotation. After washed with 1× PBS for 3 times, cells were post-fixed in 1% acrylamide (AAm, Sigma-Aldrich, A9099) and 0.7% PFA at 37°C with shaking for 4 hours and washed with 1× PBS twice for 10 min at RT. Then samples were rinsed with gelling solution consisted of 10% AAm, 0.1% N, N’-(1,2-dihydroxyethylene) bisacrylamide (DHEBA, Sigma-Aldrich, 294381), and 19% sodium acrylate (SA, Sigma-Aldrich, 408220) and 1× PBS. 0.01% ammonium persulfate (APS, Sigma-Aldrich, A3678) and 0.01% N, N, N’, N’-tetramethylethylenediamine (TEMED, Sigma-Aldrich, T22500) were added immediately before use to activate the gelling solution. Activated gelling solution was applied to samples, incubated at RT for 15 min and at 37 °C for 1 hour. Then the hydrogel was isolated and incubated in denaturation buffer (50 mM Tris-HCl, pH 6.8, 200 mM NaCl, 200 mM SDS) for 1 hour at 76 °C with rotation. After denaturation, the hydrogel was transferred to ultrapure water for two 30 min washes, and then left to expand overnight. Before image acquisition, the hydrogel was incubated in ultrapure water containing 1 μg/mL Hoechst 33342 at RT for 15 min. Then the hydrogel was put on 40 mm glass bottom dish coated with poly-L-lysine and imaged.

### Protein purification

Plasmids of pET28a-6×His-mScarlet-NuMA, pET28a-GST-NuMA and NuMA’s truncations were transferred into BL21(DE3) pLysS chemically competent cell (TransGene, CD701). Single colony was picked and cultured in Luria–Bertani (LB) medium supplemented with 30 mg/mL kanamycin at 37 °C overnight with shaking at 220 rpm. The culture was inoculated into fresh LB medium at the ratio of 1:50 to expand the culture to OD 0.6, then cooled to 18 °C. 0.5 mM IPTG was added for approximately 18 hours to induce protein expression. Cells were harvested by centrifugation at 5000 rpm, for 20 minutes 4 °C and resuspended with resuspension buffer (Tris-HCl 50 mM, pH 7.5, NaCl 2 M, glycerol 10%, 0.1 mg/mL PMSF, and 1 mM DTT for GST-tagged proteins). All purification steps were performed on ice or at 4 °C to maintain protein activity. Cell suspension was sonicated on ice, and the lysates were treated with 0.25 % PEI to eliminate DNA contamination and centrifuged at 18,000 g for 20 minutes at 4 °C. The supernatant was incubated with Ni-NTA (for proteins with 6×His tag) or GST agarose resin (for proteins with GST tag) at 4 °C for 1 hour. Ni-NTA beads were washed extensively with His washing buffer (Tris-HCl 50 mM, pH 7.5, 500 mM NaCl, imidazole 50 mM, glycerol 10%), and recombinant proteins were eluted with His elution buffer (Tris-HCl 50 mM, pH 7.5, 300 mM NaCl, imidazole 200 mM, glycerol 10%). GST agarose beads were washed extensively with GST washing buffer (Tris-HCl 50 mM, pH 7.5, 500 mM NaCl, glycerol 10%, 1 mM DTT) and recombinant proteins were eluted with GST washing buffer supplemented with 20 mM GSH.

For purification of histone H1’s isoforms, similar procedures were performed as 6×His-tagged NuMA. Plasmids of pET28a-6×His-HA-H1 were transferred into BL21(DE3) pLysS chemically competent cell and the protein expression was induced by 0.5 mM IPTG at 18 °C for approximately 18 hours. The protein was purified using Ni-NTA followed by CM Sepharose. The CM beads were extensively washed by GST washing buffer and histone H1 was eluted with CM elution buffer (Tris-HCl 50 mM, pH 7.5, 800 mM NaCl, glycerol 10%). All eluted fractions were analyzed by SDS-PAGE and Coomassie staining. Purified proteins were concentrated by ultra-centrifugation and exchanged via dialysis into storage buffer (Tris-HCl 50 mM, pH 7.5, 150 mM NaCl), flash-frozen in liquid nitrogen, sub-packaged and stored at -80 °C.

### Co-IP and pull-down

For immunoprecipitation, HEK293 cells transfected with expression plasmids or HCT116 cells with affinity tags knocked-in at NuMA gene locus were harvested by centrifugation at 200 *g*, 4 °C for 3 minutes. Cell pellets were directly lysed with IP lysis buffer (Tris-HCl 10 mM, pH 7.5, NaCl 150 mM, TritonX-100 1%, sodium deoxycholate 1%, EDTA 1mM) supplemented with 0.1 mg/mL PMSF, complete protease inhibitor and UltraNuclease on ice for 15 minutes. After lysis, the lysates were centrifuged for 10 minutes at 21,100 *g* and 4 °C. 1% of the supernatant was stored as input and the other was incubated with HA or streptavidin beads and shaking at 4 °C overnight. The next day, beads were washed by IP washing buffer (Tris-HCl 10 mM, pH 7.5, NaCl 500 mM, Tween-20 0.5%) for five times. Finally, beads with immunoprecipitated proteins were eluted with SDS-PAGE loading buffer and analyzed with western-blotting.

Purified GST or GST-NuMA and NuMA’s truncations recombinant proteins were bound to GST-magnetic beads for 2 hours at 4°C in binding buffer (Tris-HCl 10 mM, pH 7.5, NaCl 300 mM, Tween-20 0.1%, 10% glycerol), washed once with binding buffer and incubated with purified HA-tagged histone H1 overnight at 4°C with slow rotation. The next day, beads were washed three five with washing buffer (Tris-HCl 10 mM, pH 7.5, NaCl 500 mM, Tween-20 0.5%, 10% glycerol). Finally, pull-downed proteins were eluted with elution buffer (Glycine 0.1M, pH 2.5) and added with SDS-PAGE loading buffer supplemented with 50 mM Tris-HCl, pH8.8. Then samples were analyzed with western-blotting and SDS-PAGE.

For pulldown of NuMA’s C-terminal domain using immobilized nucleosomes, the mono-nucleosomes and NuMA-C were incubated with streptavidin agarose beads with or without histone H1 in the binding buffer (Tris–HCl 10 mM, pH 7.5, NaCl 150 mM, EDTA 0.2 mM, glycerol 10%, NP-40 0.1%) at 4 °C overnight. After incubation, the bound proteins were washed five times with the washing buffer (Tris–HCl 10 mM, pH 7.5, NaCl 300 mM, EDTA 0.2 mM, glycerol 10%, NP-40 0.5%, PMSF 0.5 mM) and were analyzed by Western blotting.

### Electrophoretic mobility shift assay (EMSA)

The DNA substrates used in this study were 5’-HEX labeled and resolved in reaction buffer (Tris-HCl 10 mM, pH 7.5, NaCl 150 mM). DNA substrates was commercially ordered single-strands were premixed and annealed to double-strand substrate. The DNA sequences are listed in Supplemented Table 2. Purified Histone H1 and NuMA’s truncations were mixed with dsDNA substrates and incubate at RT for 15 minutes. Then samples were added with 5% glycerol and loaded to 1% agarose gel and run electrophoresis at 80V for 1 hour in 1×TBE buffer (Tris-HCl 90 mM, pH 8.0, Boric acid 90 mM, EDTA 2 mM). HEX labeled DNA was visualized using Bio-Rad GelDOC XR.

### Nucleosome assembly and sucrose density gradient centrifugation

Mono-nucleosome, 4× and 12× nucleosome array (NA) were assembled as previously described^40^. Binding of NuMA’s C-terminal domain with H1 and 12× NA at a ratio of 1:2 was incubated in centrifugation buffer (HEPES 10 mM, pH 8.0, EDTA 0.2 mM, 25 mM NaCl) at RT for 30 minutes. The sucrose density gradient was set to 10-30 % using Gradient Master Model (Biocomp, 108). The centrifugation was performed on Beckman Coulter XL-I using a sw-60Ti rotor at 40,000× g and 4 °C under a vacuum for 16 hours. After the centrifugation was accomplished, the samples were divided into 17 fractions at same volume and then resolved on 1% agarose gel which was stained with EB (Thermo Fisher, 15585011).

### Atomic force microscopy (AFM)

NuMA’s C-terminal domain and histone H1 were incubated with 12× NA at RT for 15 minutes in AFM buffer (HEPES 20 mM, pH 7.5, 150 mM NaCl) then fixed with 0.1% glutaraldehyde on ice for 30 min. During the incubation, 20 μL spermidine (1 mM, Sigma-Aldrich, S2626) was dropped onto newly cleaved mica surface and incubate for 10 min at RT. Then rinse the mica with 200 μL ddH_2_O for 4 times and blow dry mica surface briefly. Next, the samples were diluted to 1 ng/μL and 10 μL sample solution were added onto mica surface, incubate for 10 min. Finally, wash the mica with 200 μL ddH_2_O for 3 times and blow dry gently. The prepared AFM samples were examined using Bio Atomic Force Microscope (BioScope Resolve, Bruker) and images were post-processed using Nanoscope Analysis (Bruker).

### Cryo-focused ion beam (FIB) milling and cryo-electron tomography data collection and processing

U2OS cells were seeded on grids (QUANTIFOIL SiO_2_) in DMEM medium 8 hours after transfection. Excess medium was removed by manually blotting with filter paper from the back of the grids. 4 µL of cryo buffer (cell medium with 8% v/v final concentration glycerol) was added to the grid before plunging to avoid nucleus unvitrification. Grids were blotted using a Vitrobot Mark IV at 90% humidity and 37 °C, with a blot force of 10 and a blot time of 10 seconds. Cells were plunge-frozen in an ethane/propane mixture cooled to liquid nitrogen temperatures. Grids were clipped into homemade cut-off autogrid support rings and stored in liquid nitrogen to facilitate downstream handling. Grids were loaded into an Aquilos2 cryo-focused ion beam/scanning electron microscope (FIB/SEM dual-beam microscope, Thermo Fisher Scientific) maintained at liquid nitrogen temperature throughout the procedure. To improve sample conductivity and reduce curtaining artifacts during FIB milling, grids were first sputter-coated with platinum and then coated with organometallic platinum using the gas injection system. More than 10 target cells from each strain were chosen randomly. Lamellae were prepared using a gallium ion beam at 30 kV and stage tilt angles of 13°. Rough milling was performed with currents of 0.3 nA until lamellae thickness reached ∼800 nm. Fine milling of the lamellae was gradually reduced to lower currents, with 30 pA used for the final polishing step.

Tilt series were collected using a Titan Krios G4 instrument at 300 kV, equipped with a Selectris X energy filter and a Falcon 4i camera (Thermo Fisher Scientific). Tilt series were recorded using the Tomo5 software package (Thermo Fisher Scientific). A dose-symmetric tilt scheme was used with an angular increment of 3°. Magnification was set to 64,000×, with a pixel size of 1.89 Å and a dose of 3.5 e−/Å² per tilt, for a total dose of ∼140 e-/Å². The target defocus ranged from -4.0 to -5.0 µm. Data were processed using the TOMOMAN^75^ package version 0.7 pipeline (https://github.com/williamnwan/TOMOMAN). After frame motion correction, bad tilts were removed after manual inspection using a TOMOMAN script. Tilt series were split into odd and even tilts during motion correction for denoising, and the resulting stacks were denoised using CryoCARE^76^. Tilt series were aligned using AreTomo^77^ version 1.3.3. Tomogram reconstructions were performed with IMOD^78^ version 4.12.32 at bin4 for visualization and bin2 for nucleosome template matching. Nucleosome coordinates were determined using the template matching routine from STOPGAP^79^ version 0.7. A reference nucleosome core particle density map (EMD-8140)^80^ was lowpass filtered to 30 Å to serve as the template. Subtomograms were binned by a factor of 2 (pixel size 3.78 Å/pixel), extracted by Warp^81^ from best quality 15 tomograms (10 tomograms from NuMA-dC, 5 tomograms from NuMA-FL). After 3D classification in RELION 3.0^82^ with spherical mask, 4709 particles from NuMA-dC and 3601 particles from NuMA-FL reasonable classes were retained for auto-refinement. The half maps and particle lists were imported into M^83^ for geometric refinement. Bin1 (pixel size 1.89 Å/pixel) subtomograms were re-extracted after M^83^ refinement and imported into RELION^82^ for a second round of 3D classification. Relative particles were retained for M geometric and CTF parameter refinement. After reaching resolution convergence, a final map at 9.9 Å resolution from dC cells and 17.8 Å from FL cells were obtained within the reconstruction mask using the 0.143 FSC criterion.

NND analysis was based on nucleosome coordinates after subtomogram alignment. Nucleosome core particle (PDB:2CV5)^84^ and chromatosome (PDB: 7PF6)^85^ were applied for EM map docking.

## Data availability

The sequencing data of Hi-C, ATAC-seq, RNA-seq and Cut&Tag analyzed in this study are accessible at the NCBI GEO website under the accession number GSE227941. R markdown scripts enabling the main analyses are available from the responding author upon reasonable request. EM density maps have been deposited to EMDB under codes EMD-51858, EMD-51859. All other data supporting the findings of this study are available from the corresponding author on reasonable request.

## Code availability

Code use in this article can be made available upon request to the corresponding author.

## Acknowledgments

We thank Prof. Dongyi Xu for providing HCT116 cells. We thank National Center for Protein Science at Peking University in Beijing, China, for assistance with flow cytometry and imaging, particularly Ms. Liying Du, Dr. Chunyan Shan, Dr. Liqin Fu and Dr. Yiqun Liu for technical help. This work is supported by grants from the National Key R&D Program of China No. 2022YFA1303103 and 2022YFA3401100, and the National Science Foundation of China 22127804 for Y.S.

## Author information

These authors contributed equally: Yao Wang, Wenxue Zhao, Jiahao Niu.

## Contributions

Y.S. and Y.W. conceived and designed this study. Y.W. and J.N. performed most biochemistry and cell biology experiments. W.Z. performed sequencing library construction and bioinformatics analysis. C. L. performed nucleosomes array reconstitution. X.W. performed super-resolution imaging quantification and analysis. P.X. performed cryo-electron tomography and data processing. W.Y. performed AFM. S. A. provided assistance with CUT&Tag experiments. Y.W., W.Z., J.N. and Y.S. wrote this paper. All authors participated in the discussion of the manuscript, and the manuscript was written through contributions of all authors. All authors have given approval to the final version of the manuscript.

## References

1. Grewal, S.I. & Jia, S. Heterochromatin revisited. Nat Rev Genet 8, 35–46 (2007).

2. Allshire, R.C. & Madhani, H.D. Ten principles of heterochromatin formation and function. Nat Rev Mol Cell Biol 19, 229–244 (2018).

3. Janssen, A., Colmenares, S.U. & Karpen, G.H. Heterochromatin: Guardian of the Genome. Annu Rev Cell Dev Biol 34, 265–288 (2018).

4. Greenstein, R.A. & Al-Sady, B. Epigenetic fates of gene silencing established by heterochromatin spreading in cell identity and genome stability. Curr Genet 65, 423–428 (2019).

5. Grewal, S.I.S. The molecular basis of heterochromatin assembly and epigenetic inheritance. Mol Cell 83, 1767–1785 (2023).

6. Fan, H. et al. The nuclear matrix protein HNRNPU maintains 3D genome architecture globally in mouse hepatocytes. Genome Res 28, 192–202 (2018).

7. Nozawa, R.S. et al. SAF-A Regulates Interphase Chromosome Structure through Oligomerization with Chromatin-Associated RNAs. Cell 169, 1214–1227 e1218 (2017).

8. Ma, G. et al. The nuclear matrix stabilizes primed-specific genes in human pluripotent stem cells. Nat Cell Biol 27, 232–245 (2025).

9. Chang, L. et al. Nuclear peripheral chromatin-lamin B1 interaction is required for global integrity of chromatin architecture and dynamics in human cells. Protein Cell 13, 258–280 (2020).

10. Huo, X. et al. The Nuclear Matrix Protein SAFB Cooperates with Major Satellite RNAs to Stabilize Heterochromatin Architecture Partially through Phase Separation. Mol Cell 77, 368–383 e367 (2020).

11. Zhang, Y. et al. MATR3-antisense LINE1 RNA meshwork scaffolds higher-order chromatin organization. EMBO Rep 24, e57550 (2023).

12. Berezney, R. & Coffey, D.S. Nuclear matrix. Isolation and characterization of a framework structure from rat liver nuclei. J Cell Biol 73, 616–637 (1977).

13. Harborth, J., Wang, J., Gueth-Hallonet, C., Weber, K. & Osborn, M. Self assembly of NuMA: multiarm oligomers as structural units of a nuclear lattice. EMBO J 18, 1689–1700 (1999).

14. Zeng, C., He, D. & Brinkley, B.R. Localization of NuMA protein isoforms in the nuclear matrix of mammalian cells. Cell Motil Cytoskeleton 29, 167–176 (1994).

15. Engelke, R. et al. The quantitative nuclear matrix proteome as a biochemical snapshot of nuclear organization. J Proteome Res 13, 3940–3956 (2014).

16. Lydersen, B.K. & Pettijohn, D.E. Human-specific nuclear protein that associates with the polar region of the mitotic apparatus: distribution in a human/hamster hybrid cell. Cell 22, 489–499 (1980).

17. Compton, D.A., Szilak, I. & Cleveland, D.W. Primary Structure of NuMA, an Intranuclear Protein That Defines a Novel Pathway for Segregation of Proteins at Mitosis. Journal of Cell Biology 116, 1395–1408 (1992).

18. Kempf, T., Bischoff, F.R., Kalies, I. & Ponstingl, H. Isolation of Human NuMA Protein. Febs Lett 354, 307–310 (1994).

19. Radulescu, A.E. & Cleveland, D.W. NuMA after 30 years: the matrix revisited. Trends Cell Biol 20, 214–222 (2010).

20. Kiyomitsu, T. & Boerner, S. The Nuclear Mitotic Apparatus (NuMA) Protein: A Key Player for Nuclear Formation, Spindle Assembly, and Spindle Positioning. Front Cell Dev Biol 9, 653801 (2021).

21. Luderus, M.E., den Blaauwen, J.L., de Smit, O.J., Compton, D.A. & van Driel, R. Binding of matrix attachment regions to lamin polymers involves single-stranded regions and the minor groove. Mol Cell Biol 14, 6297–6305 (1994).

22. Rajeevan, A., Keshri, R., Kapoor, S. & Kotak, S. NuMA interaction with chromatin is vital for proper chromosome decondensation at the mitotic exit. Mol Biol Cell 31, 2437–2451 (2020).

23. Serra-Marques, A. et al. The mitotic protein NuMA plays a spindle-independent role in nuclear formation and mechanics. J Cell Biol 219, 1–18 (2020).

24. Kivinen, K., Taimen, P. & Kallajoki, M. Silencing of Nuclear Mitotic Apparatus protein (NuMA) accelerates the apoptotic disintegration of the nucleus. Apoptosis 15, 936–945 (2010).

25. Abad, P.C. et al. NuMA influences higher order chromatin organization in human mammary epithelium. Mol Biol Cell 18, 348–361 (2007).

26. Gueth-Hallonet, C., Wang, J., Harborth, J., Weber, K. & Osborn, M. Induction of a regular nuclear lattice by overexpression of NuMA. Exp Cell Res 243, 434–452 (1998).

27. Hergeth, S.P. & Schneider, R. The H1 linker histones: multifunctional proteins beyond the nucleosomal core particle. EMBO Rep 16, 1439–1453 (2015).

28. Natsume, T., Kiyomitsu, T., Saga, Y. & Kanemaki, M.T. Rapid Protein Depletion in Human Cells by Auxin-Inducible Degron Tagging with Short Homology Donors. Cell Rep 15, 210–218 (2016).

29. Silk, A.D., Holland, A.J. & Cleveland, D.W. Requirements for NuMA in maintenance and establishment of mammalian spindle poles. J Cell Biol 184, 677–690 (2009).

30. Rao, S.S. et al. A 3D map of the human genome at kilobase resolution reveals principles of chromatin looping. Cell 159, 1665–1680 (2014).

31. Buenrostro, J.D., Giresi, P.G., Zaba, L.C., Chang, H.Y. & Greenleaf, W.J. Transposition of native chromatin for fast and sensitive epigenomic profiling of open chromatin, DNA-binding proteins and nucleosome position. Nat Methods 10, 1213–1218 (2013).

32. Ernst, J. & Kellis, M. ChromHMM: automating chromatin-state discovery and characterization. Nat Methods 9, 215–216 (2012).

33. Willcockson, M.A. et al. H1 histones control the epigenetic landscape by local chromatin compaction. Nature 589, 293–298 (2021).

34. Ricci, M.A., Manzo, C., Garcia-Parajo, M.F., Lakadamyali, M. & Cosma, M.P. Chromatin fibers are formed by heterogeneous groups of nucleosomes in vivo. Cell 160, 1145–1158 (2015).

35. Fyodorov, D.V., Zhou, B.R., Skoultchi, A.I. & Bai, Y. Emerging roles of linker histones in regulating chromatin structure and function. Nat Rev Mol Cell Biol 19, 192–206 (2018).

36. Woodcock, C.L., Skoultchi, A.I. & Fan, Y. Role of linker histone in chromatin structure and function: H1 stoichiometry and nucleosome repeat length. Chromosome Res 14, 17–25 (2006).

37. Baldi, S., Korber, P. & Becker, P.B. Beads on a string-nucleosome array arrangements and folding of the chromatin fiber. Nat Struct Mol Biol 27, 109–118 (2020).

38. Kaya-Okur, H.S., Janssens, D.H., Henikoff, J.G., Ahmad, K. & Henikoff, S. Efficient low-cost chromatin profiling with CUT&Tag. Nature Protocols 15, 3264–3283 (2020).

39. Hizume, K., Yoshimura, S.H. & Takeyasu, K. Linker histone H1 per se can induce three-dimensional folding of chromatin fiber. Biochemistry 44, 12978–12989 (2005).

40. Song, F. et al. Cryo-EM study of the chromatin fiber reveals a double helix twisted by tetranucleosomal units. Science 344, 376–380 (2014).

41. Pownall, M.E. et al. Chromatin expansion microscopy reveals nanoscale organization of transcription and chromatin. Science 381, 92–99 (2023).

42. M’Saad, O. & Bewersdorf, J. Light microscopy of proteins in their ultrastructural context. Nature Communications 11 (2020).

43. Lu, X. et al. Drosophila H1 regulates the genetic activity of heterochromatin by recruitment of Su(var)3-9. Science 340, 78–81 (2013).

44. Liu, C.F. et al. Histone H1 facilitates restoration of H3K27me3 during DNA replication by chromatin compaction. Nature Communications 14 (2023).

45. Healton, S.E. et al. H1 linker histones silence repetitive elements by promoting both histone H3K9 methylation and chromatin compaction. Proc Natl Acad Sci U S A 117, 14251–14258 (2020).

46. He, S. et al. Linker histone H1 drives heterochromatin condensation via phase separation in Arabidopsis. Plant Cell (2024).

47. Lieberman-Aiden, E. et al. Comprehensive mapping of long-range interactions reveals folding principles of the human genome. Science 326, 289–293 (2009).

48. Gibson, B.A. et al. Organization of Chromatin by Intrinsic and Regulated Phase Separation. Cell 179, 470–484 e421 (2019).

49. Lu, X. et al. Linker histone H1 is essential for Drosophila development, the establishment of pericentric heterochromatin, and a normal polytene chromosome structure. Genes Dev 23, 452–465 (2009).

50. Hayward, A. & Gilbert, C. Transposable elements. Current Biology 32, R904–R909 (2022).

51. Wells, J.N. & Feschotte, C. A Field Guide to Eukaryotic Transposable Elements. Annual Review of Genetics, Vol 54, 2020 54, 539–561 (2020).

52. Xu, R.M. et al. Stage-specific H3K9me3 occupancy ensures retrotransposon silencing in human pre-implantation embryos. Cell Stem Cell 29, 1051-+ (2022).

53. Liu, X.Q. et al. Resurrection of endogenous retroviruses during aging reinforces senescence. Cell 186, 287-+ (2023).

54. Yan, L. et al. Single-cell RNA-Seq profiling of human preimplantation embryos and embryonic stem cells. Nat Struct Mol Biol 20, 1131–1139 (2013).

55. Wu, J. et al. Chromatin analysis in human early development reveals epigenetic transition during ZGA. Nature 557, 256–260 (2018).

56. Deng, Q., Ramsköld, D., Reinius, B. & Sandberg, R. Single-cell RNA-seq reveals dynamic, random monoallelic gene expression in mammalian cells. Science 343, 193–196 (2014).

57. Turner, A.L. et al. Highly disordered histone H1-DNA model complexes and their condensates. Proc Natl Acad Sci U S A 115, 11964–11969 (2018).

58. Wang, L. et al. Rett syndrome-causing mutations compromise MeCP2-mediated liquid-liquid phase separation of chromatin. Cell Res 30, 393–407 (2020).

59. Sun, M. et al. NuMA regulates mitotic spindle assembly, structural dynamics and function via phase separation. Nature Communications 12 (2021).

60. Ma, H. et al. NuMA forms condensates through phase separation to drive spindle pole assembly. Journal of Molecular Cell Biology 14 (2022).

61. Okumura, M., Natsume, T., Kanemaki, M.T. & Kiyomitsu, T. Dynein-Dynactin-NuMA clusters generate cortical spindle-pulling forces as a multi-arm ensemble. Elife 7, 1–24 (2018).

62. Jackson, D.A. & Pombo, A. Replicon clusters are stable units of chromosome structure: Evidence that nuclear organization contributes to the efficient activation and propagation of S phase in human cells. Journal of Cell Biology 140, 1285–1295 (1998).

63. Dimitrova, D.S., Todorov, I.T., Melendy, T. & Gilbert, D.M. Mcm2, but not RPA, is a component of the mammalian early G1-phase prereplication complex. J Cell Biol 146, 709–722 (1999).

64. Servant, N. et al. HiC-Pro: an optimized and flexible pipeline for Hi-C data processing. Genome Biol. 16, 259 (2015).

65. Servant, N. et al. HiTC: exploration of high-throughput ‘C’ experiments. Bioinformatics 28, 2843–2844 (2012).

66. Durand, N.C. et al. Juicer Provides a One-Click System for Analyzing Loop-Resolution Hi-C Experiments. Cell Syst 3, 95–98 (2016).

67. Langmead, B. & Salzberg, S.L. Fast gapped-read alignment with Bowtie 2. Nat Methods 9, 357–359 (2012).

68. Ramirez, F. et al. deepTools2: a next generation web server for deep-sequencing data analysis. Nucleic Acids Res 44, W160–165 (2016).

69. Zhang, Y. et al. Model-based analysis of ChIP-Seq (MACS). Genome Biol 9, R137 (2008).

70. Quinlan, A.R. BEDTools: The Swiss-Army Tool for Genome Feature Analysis. Curr Protoc Bioinformatics 47, 11 12 11–34 (2014).

71. Robinson, J.T., Thorvaldsdottir, H., Turner, D. & Mesirov, J.P. igv.js: an embeddable JavaScript implementation of the Integrative Genomics Viewer (IGV). Bioinformatics 39 (2023).

72. Anders, S., Pyl, P.T. & Huber, W. HTSeq--a Python framework to work with high-throughput sequencing data. Bioinformatics 31, 166–169 (2015).

73. Love, M.I., Huber, W. & Anders, S. Moderated estimation of fold change and dispersion for RNA-seq data with DESeq2. Genome Biol 15, 550 (2014).

74. Kuleshov, M.V. et al. Enrichr: a comprehensive gene set enrichment analysis web server 2016 update. Nucleic Acids Res 44, W90–97 (2016).

75. Khavnekar, S., Erdmann, P. & Wan, W. TOMOMAN: Streamlining Cryo-electron Tomography and Subtomogram Averaging Workflows Using TOMOgram MANager. Microscopy and Microanalysis 29, 1020–1020 (2023).

76. Buchholz, T.-O., Jordan, M., Pigino, G. & Jug, F. in 2019 IEEE 16th International Symposium on Biomedical Imaging (ISBI 2019) 502–506 (2019).

77. Zheng, S., et al. AreTomo: An integrated software package for automated marker-free, motion-corrected cryo-electron tomographic alignment and reconstruction. Journal of Structural Biology: X 6 (2022).

78. Kremer, J.R., Mastronarde, D.N. & McIntosh, J.R. Computer Visualization of Three-Dimensional Image Data Using IMOD. Journal of Structural Biology 116, 71–76 (1996).

79. Wan, W., Khavnekar, S. & Wagner, J. STOPGAP: an open-source package for template matching, subtomogram alignment and classification. Acta Crystallographica Section D Structural Biology 80, 336–349 (2024).

80. Chua, E.Y.D. et al. 3.9 Å structure of the nucleosome core particle determined by phase-plate cryo-EM. Nucleic Acids Research 44, 8013–8019 (2016).

81. Tegunov, D. & Cramer, P. Real-time cryo-electron microscopy data preprocessing with Warp. Nat Meth 16, 1146–1152 (2019).

82. Zivanov, J. et al. New tools for automated high-resolution cryo-EM structure determination in RELION-3. eLife 7 (2018).

83. Tegunov, D., Xue, L., Dienemann, C., Cramer, P. & Mahamid, J. Multi-particle cryo-EM refinement with M visualizes ribosome-antibiotic complex at 3.5 Å in cells. Nat Meth 18, 186–193 (2021).

84. Tsunaka, Y. Alteration of the nucleosomal DNA path in the crystal structure of a human nucleosome core particle. Nucleic Acids Research 33, 3424–3434 (2005).

85. Dombrowski, M., Engeholm, M., Dienemann, C., Dodonova, S. & Cramer, P. Histone H1 binding to nucleosome arrays depends on linker DNA length and trajectory. Nature Structural & Molecular Biology 29, 493–501 (2022).

