## Supplementary figures and tables for "NuMA promotes constitutive heterochromatin compaction by stabilizing linker histone H1 on chromatin"

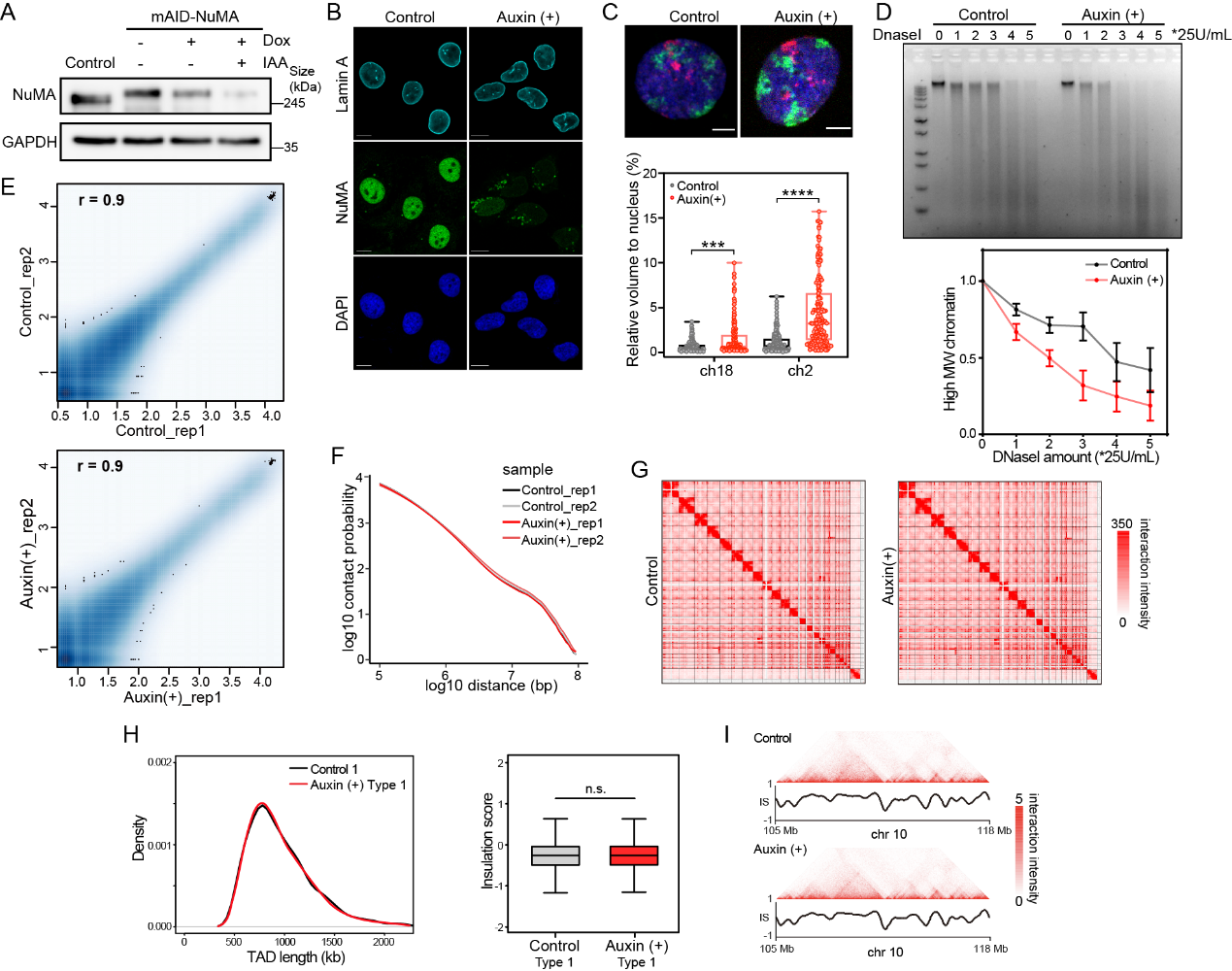

#### Figure S1. NuMA regulates global 3D genome organization

A. Schematic of auxin-inducible degron system (upper) and NuMA-depletion induced by auxin in unmodified U2OS cells as control and U2OS-mAID-NuMA cells detected by western blotting (lower).
B. IF imaging of Lamin A in untreated U2OS-mAID-NuMA cells as control and NuMA-depleted U2OS -mAID-NuMA cells induced by auxin. Scale bar, 10 μm.
C. FISH imaging (upper) and quantification (lower) of the volumes occupied by chromosome 2 (green) and 18 (red) relative to the nucleus volume in untreated U2OS-mAID-NuMA cells as control and NuMA-depleted U2OS-mAID-NuMA cells induced by auxin. Error bars represent SD (n≥50). ****p < 0.0005, Mann-Whitney test. Scale bar, 5 μm.
D. Global chromatin accessibility upon NuMA-depletion detected by DNaseI digestion assay, with untreated U2OS-mAID-NuMA cells as control. Agarose gel image of genomic DNA digested by DNaseI at different concentrations (left), and percentages of high-molecular-weight (MW) genomic DNA (>5 kb) (right). The gel is representative of three independent experiments. Error bars represent SD (n=3).
E. Pearson correlation coefficients of the whole genome contact matrices (resolution: 500 kb) of two replicates in the control and NuMA-depleted HCT116-mAID-NuMA cells induced by auxin. Duplicate Hi-C libraries were generated for each condition, with correlation analysis demonstrating a high degree of consistency among replicates.
F. Hi-C interaction frequency as a function of genomic linear distance for control and NuMA-depletion replicates.
G. Hi-C contact maps of control and auxin-induced HCT116-mAID-NuMA cells at 500 kb resolution. All matrices were normalized to the same sequencing depth.
H. TAD length distribution and insulation score of Type 1 chromatin.
I. Example of TAD pattern and insulation score distribution for chromosome 10 (105-118 Mb) in control and NuMA-depleted HCT116-mAID-NuMA cells.

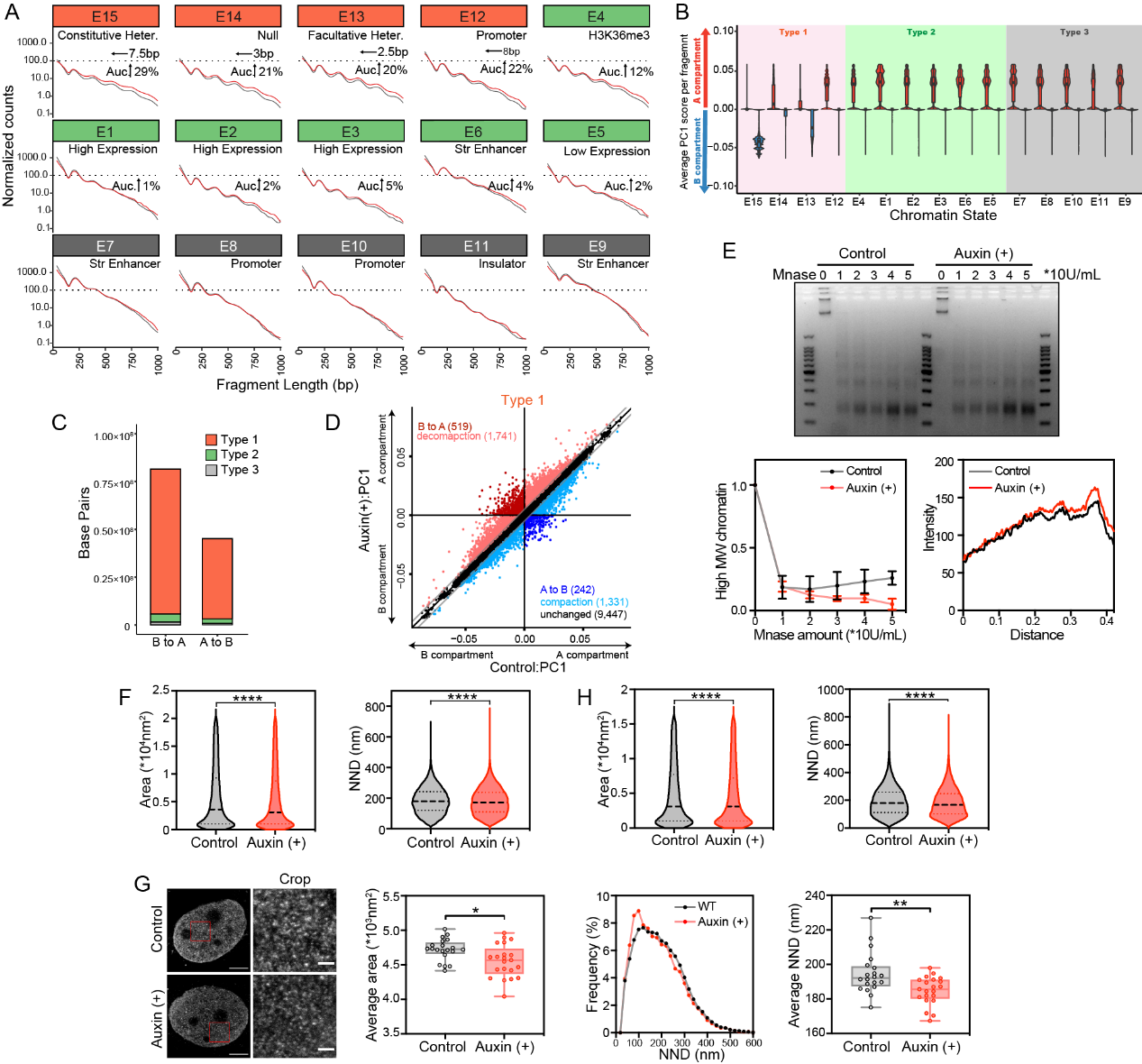

#### Figure S2. NuMA regulates heterochromatin compaction by maintaining nucleosome stacking

A. Changes in NRL and chromatin accessibility of 15 chromatin states in untreated HCT116-mAID-NuMA cells as control and NuMA-depleted HCT116-mAID-NuMA cells induced by auxin. Heter, heterochromatin; Str Enhanc, strong enhancer.
B. The PC1 score for each genomic region grouped by 15 chromatin states and Type 1, 2 and 3.
C. Proportion of Type 1, 2, and 3 chromatin in compartment A/B switch regions of NuMA-depleted HCT116-mAID-NuMA cells.
D. Scatterplot showing compartment A/B switches in Type 1 chromatin after NuMA-depletion. PC1 was calculated for each 150 kb genomic segment to define A and B compartments and identify compartment switching, decompaction and compaction upon NuMA-depletion.
E. Chromatin accessibility upon NuMA-depletion in U2OS-mAID-NuMA cells as detected by Mnase digestion assay. Gel image of genomic DNA digested by Mnase at different concentrations (left), percentages of MW genomic DNA (>5 kb) (middle), and an example of nucleosome ladders at the well 2 (right). Error bars represent SD (n=3).
F. Quantification of the area and NND of nucleosome clutches acquired from STED imaging of H2B in HCT116-mAID-NuMA cells. Error bars represent SD (n=20). ****p < 0.0005, Mann-Whitney test.
G. STED imaging of H2B (left) and quantification (right) of nucleosome clutches of U2OS-mAID-NuMA cells, including the average area, frequency of NND and average NND. Error bars represent SD (n=20). ****p < 0.0005, **p < 0.05, *p < 0.5, Mann-Whitney test. Scale bar, 5 μm (left) and 1 μm (right).
H. Quantification of the area and NND of nucleosome clutches acquired from STED imaging of H2B in U2OS-mAID-NuMA cells. Error bars represent SD (n=20). ****p < 0.0005, Mann-Whitney test.

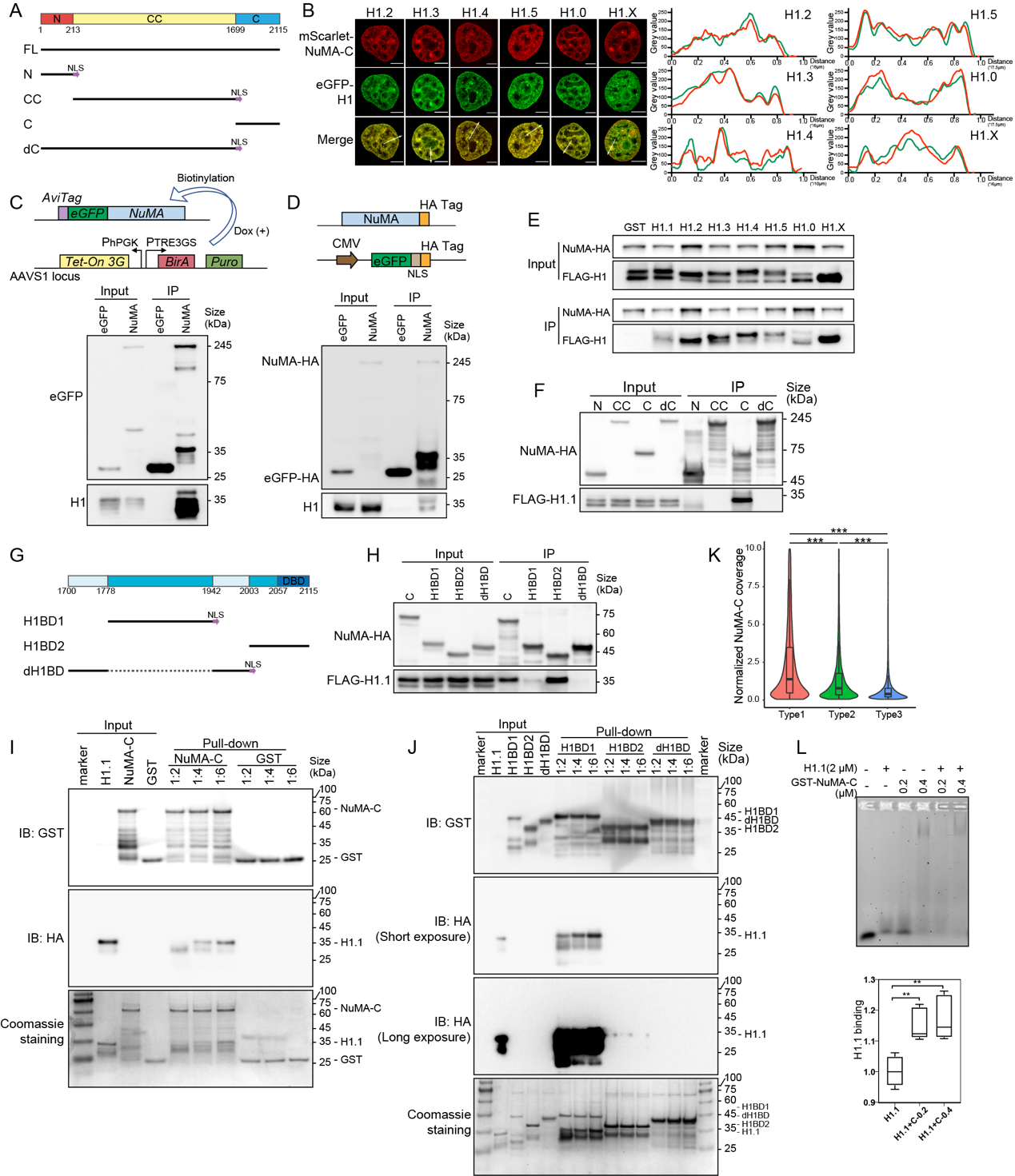

#### Figure S3. NuMA promotes linker histone H1’s binding to chromatin

A． Diagram of structural domains of and truncations of NuMA. All truncations were fused with mScarlet at the N-terminus, and nuclear localization signal (NLS) were added at the C-terminus of N, CC and dC.
B． Representative live-cell image of over-expressed NuMA truncations with H1.2, H1.3, H1.4, H1.5, H1,0 and H1.X in HeLa cells. Plots of the red and green pixel intensities along the white arrow in the left panel. Scale bar, 5 μm.
C． Scheme for specific biotinylation of NuMA in HCT116 cells (upper). An Avitag epitope and eGFP was knocked-in N-terminal of the endogenous NUMA gene. The Avitag is biotinylated by BirA, expressed from the *AAVSI* locus induced by doxycycline. Biotinylation of tagged endogenous NuMA and its immunoprecipitation with H1 in HCT116 cells shown by western-blot (lower).
D． Scheme for an HA tag knocked in the C-terminal of endogenous NuMA1 gene in HCT116 cells and over-expressed eGFP-HA served as negative control (upper). Immunoblots showing the immunoprecipitation of endogenous NuMA with HA tag and H1 of HCT116 cells (lower).
E． Immunoblots showing the immunoprecipitations of HA-tagged NuMA and FLAG-tagged histone H1.1, H1.2, H1.3, H1.4, H1.5, H1,0 and H1.X over-expressed in 293T cells.
F． Immunoblots showing the immunoprecipitation of HA-tagged NuMA truncations and FLAG-tagged H1.1 over-expressed in 293T cells.
G． Diagram of structural domains of and truncations of NuMA-C. All truncations were fused with mScarlet at the N-terminus, and NLS were added at the C-terminus of H1BD1 and dH1BD.
H． Immunoblots showing the immunoprecipitation of HA-tagged truncations of NuMA’s C-terminal and FLAG-tagged H1.1 over-expressed in 293T cells.
I． Pull-down showing the immunoprecipitation of NuMA-C and H1.1 purified from E. coli.
J． Pull-down showing the immunoprecipitation of truncations of NuMA-C and H1.1 purified from E. coli.
K． Normalized CUT&Tag signals of NuMA-C in each chromatin type.
L． The effect of increased doses of NuMA-C on H1.1 as detected by EMSA analysis (upper) and quantification of DNA bound by H1.1 and NuMA-C (lower). Error bars represent SD (n=3). **p < 0.05, Mann-Whitney test.

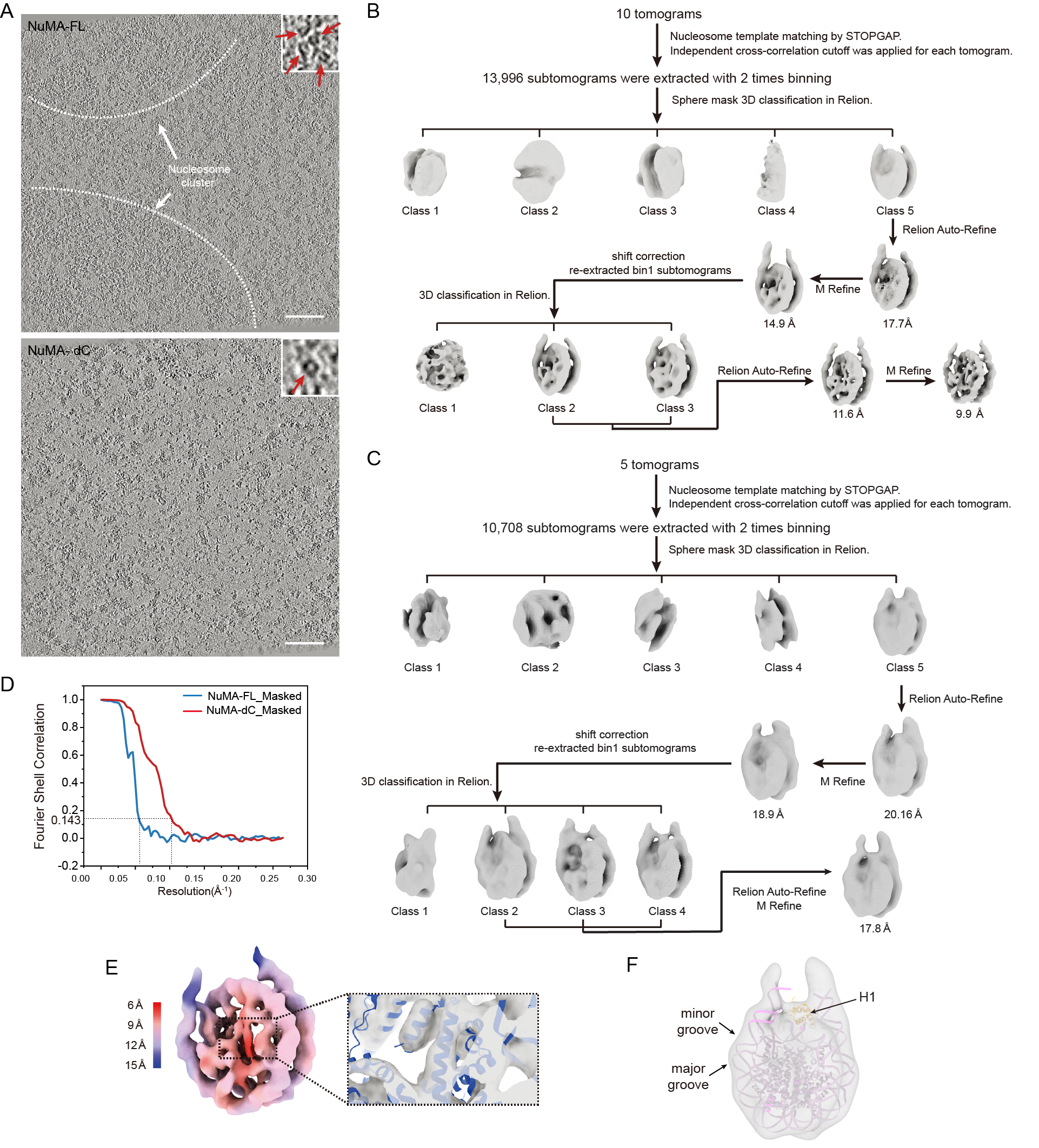

#### Figure S4. Cryo-FIB-tomography and *in situ* nucleosome subtomogram averaging workflow

A. Relative tomogram slices to Figure 4J. Dense nucleosome clusters in cells overexpressing NuMA-FL were labeled with white dot line. Zoomed in views in the upper right corners, and mono nucleosomes indicated with red arrows. Scale bar, 100 nm.
B. Subtomogram averaging workflow for cells overexpression NuMA-dC.
C. Subtomogram averaging workflow for cells overexpression NuMA-FL.
D. FSC curves of NuMA-FL and dC masked density maps.
E. *In situ* nucleosome electron map from NuMA-dC colored based on local resolution indicated on the left. Crystal structure (in blue) was fitted into the map as zoomed-in image on the right.
F. *In situ* nucleosome electron map from NuMA-FL and crystal structure of chromatosome (PDB: 7PF6) fitted into the map. H1’s global domain labeled yellow. Features labeled with black arrows.

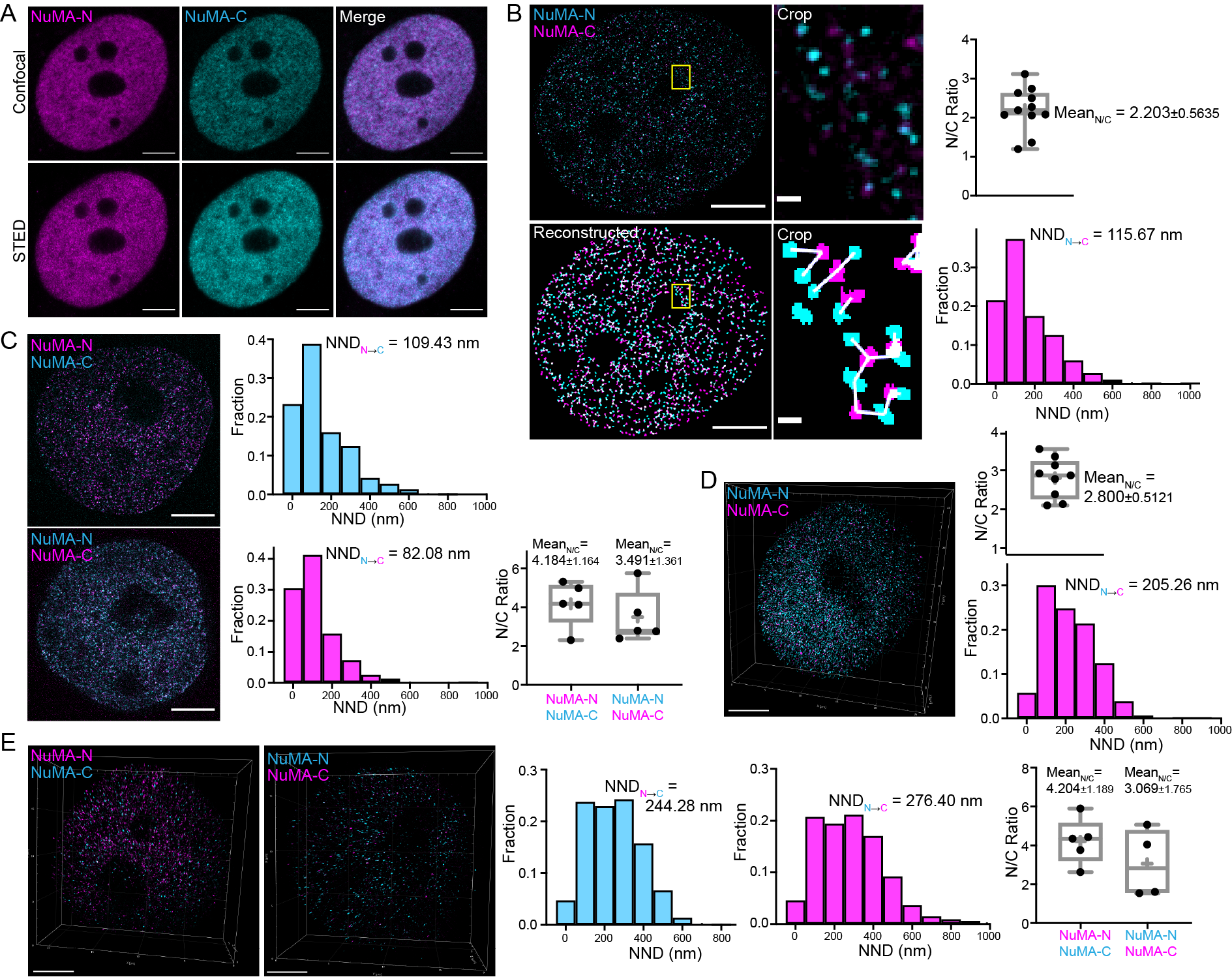

#### Figure S5. NuMA oligomerizes into quasi-network organization through its C-termini *in vivo*

A. Confocal and STED images of NuMA-N and NuMA-C in U2OS cells. Scale bar, 5 μm.
B. Expanded 2D IF images of NuMA-N and NuMA-C and reconstructed NuMA oligomers in U2OS cells (left). NuMA-N was labeled by Alexa594 and NuMA-C was labeled by Atto647N. Calculated ratio of numbers and distribution of NND from NuMA-C to NuMA-N clusters in U2OS 2D expanded IF images (right). Scale bars, 5 μm and 250 nm.
C. Expanded 2D IF images of NuMA-N and NuMA-C, reconstructed NuMA oligomers, calculated ratio of numbers of NuMA-N and NuMA-C clusters and distribution of NND from NuMA-C to NuMA-N clusters in HCT116 cells. NuMA-N was labeled by Atto647N and Alexa594 respectively, and NuMA-C was labeled by Atto647N. and Alexa594 respectively. Scale bar, 5 μm.
D. Perspective 3D view of expanded IF images of NuMA-N and NuMA-C in U2OS cells. NuMA-N was labeled by Alexa594 and NuMA-C was labeled by Atto647N. Calculated ratio of numbers and distribution of NND from NuMA-C to NuMA-N clusters in U2OS 3D expanded IF images. Scale bar, 5 μm.
E. Perspective 3D view of expanded IF images of NuMA-N and NuMA-C in HCT116 cells. NuMA-N was labeled by Atto647N and Alexa594 respectively, and NuMA-C was labeled by Atto647N. and Alexa594 respectively. Calculated ratio of numbers and distribution of NND from NuMA-C to NuMA-N clusters in HCT116 cells. Scale bar, 5 μm.

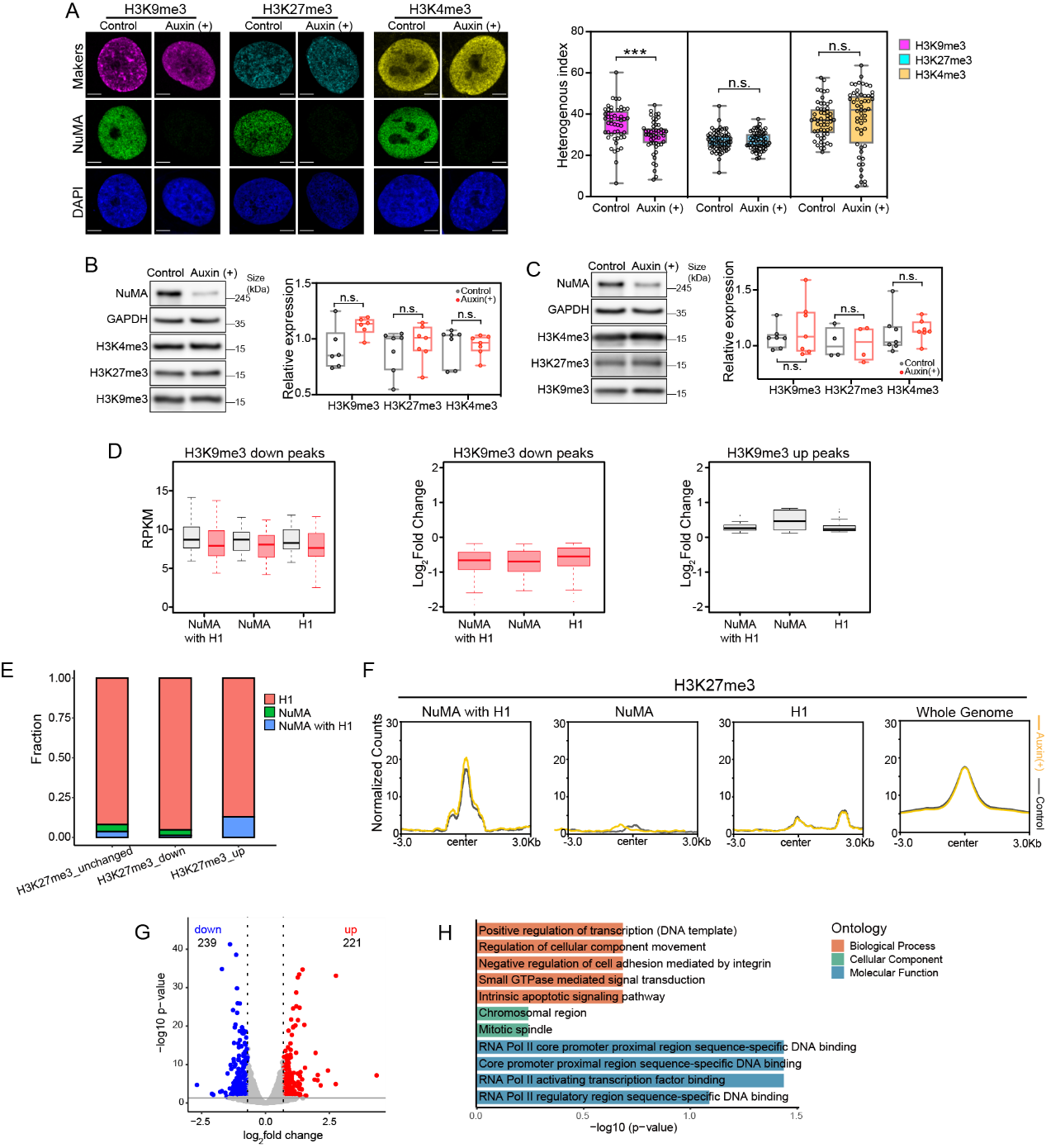

#### Figure S6. NuMA contributes to epigenetic maintenance of constitutive heterochromatin and repression of LTR expression

A. IF imaging (left) and quantification of the heterogenous index (right) of H3K9me3, H3K27me3 and H3K4me3 in U2OS-mAID-NuMA cells after induced by auxin, with untreated U2OS-mAID-NuMA cells as control. Error bars represent SD (n≥50). ****p < 0.0005, Mann-Whitney test. Scale bar, 5 μm.
B. Western blotting (left) and quantification (right) of H3K4me3, H3K27me3 and H3K9me3 expression levels after auxin treatment in HCT116-mAID-NuMA cells. Error bars represent SD (n=3).
C. Western blotting (left) and quantification (right) of H3K4me3, H3K27me3 and H3K9me3 expression levels after auxin treatment in U2OS-mAID-NuMA cells. Error bars represent SD (n=3). All gels are representative of three independent experiments.
D. Expression level of transposable elements with down-regulated H3K9me3 in genomic regions where NuMA-C and H1 are co-bound and genomic regions bound by NuMA or H1 alone (left). Expression changes of transposable elements in genomic regions with down-regulated H3K9me3 (middle) and up-regulated H3K9me3 (right) upon NuMA-depletion.
E. Fraction of NuMA-C and H1 enrichment in genomic regions with unchanged H3K27me3, down-regulated H3K27me3 and up-regulated H3K27me3 upon NuMA-depletion.
F. Change of H3K27me3 CUT&Tag peaks upon NuMA-depletion in different categories (regions where NuMA-C and H1 are co-bound and genomic regions bound by NuMA or H1 alone), with the change profile in the whole genome as control. The peaks were aligned using the center of the peaks.
G. Volcano plot showing differentially expressed genes in NuMA-depleted HCT116-mAID-NuMA cells, with untreated HCT116-mAID-NuMA cells as control. Log2 fold change cut-off, 0.7; P value cut-off, 0.05.
H. GO analysis of up-regulated genes upon NuMA-depletion.

### Supplementary Tables

#### Table S1. Antibodies

| Antibody Designation | Source or reference | Identifiers | Additional information |
| --- | --- | --- | --- |
| NuMA (Rabbit, A region within amino acids 1 and 309 of Human NuMA) | Abcam | ab97585 | 1:1000 for western blotting and 1:200 for immunofluorescence |
| NuMA (Mouse, Amino acids 1816-2115 mapping at the C-terminus of human NuMA) | Santa Cruz | sc-365532 | 1:500 for western blotting, 1:100 for immunofluorescence and 1mg/ml for CUT&Tag |
| NuMA (Rabbit, Full-length recombinant NuMA) | Invitrogen | PA1-32451 | 1:1000 for western blotting |
| NuMA (Mouse, A region within amino acids 1 and 309 of NuMA) | Invitrogen | MA5-17293 | 1:1000 for western blotting and 1:200 for immunofluorescence |
| NuMA (Rabbit, A region within amino acids 1 and 309 of NuMA) | Invitrogen | PA5-22285 | 1:1000 for western blotting and 1:200 for immunofluorescence |
| Lamin A (Mouse) | Abcam | ab8980 | 1:200 for immunofluorescence |
| GAPDH (Mouse) | Yeasen | 30201ES20 | 1:5000 for western blotting |
| H3K9me3 (Rabbit) | Abcam | ab8898 | 1:200 for immunofluorescence and 1:1000 for western blotting |
| H3K27me3 (Rabbit) | CST | 9733S | 1:200 for immunofluorescence and 1:1000 for western blotting |
| H3K4me3 (Rabbit) | CST | 9725T | 1:200 for immunofluorescence and 1:1000 for western blotting |
| FLAG (Rabbit) | Proteintech | 20543-1-AP | 1:1000 for western blotting |
| GST (Rabbit) | Yeasen | 30902ES60 | 1:1000 for western blotting |
| HA (Rabbit) | Yeasen | 30702ES60 | 1:1000 for western blotting |
| H1 (Mouse) | Santa Cruz | sc-8030 | 1:500 for western blotting and 1mg/ml for CUT&Tag |
| H2B (Rabbit) | Abcam | Ab1790 | 1:200 for immunofluorescence |
| H1 (Mouse) | Millipore | 05-457 | 1:200 for immunofluorescence and 1:1000 for western blotting |
| H3 (Rabbit) | Abcam | Ab1791 | 1:2000 for western blotting |
| Goat anti-mouse IgG (H+L)-HRP conjugated | Easybio | BE0102 | 1:5000 for western blotting |
| Goat anti-rabbit IgG (H+L)-HRP conjugated | Easybio | BE0101 | 1:10000 for western blotting |
| Donkey anti-mouse IgG (H+L) highly cross-absorbed secondary antibody, Alexa Fluor 488 | Thermo Fisher | A-21202 | 1:200 for immunofluorescence |
| Donkey anti-rabbit IgG (H+L) highly cross-absorbed secondary antibody, Alexa Fluor 488 | Thermo Fisher | A-21206 | 1:200 for immunofluorescence |
| Goat anti-mouse IgG (H+L) highly cross-absorbed secondary antibody, Alexa Fluor 594 | Thermo Fisher | A-11032 | 1:200 for immunofluorescence |
| Donkey anti-rabbit IgG (H+L) highly cross-absorbed secondary antibody, Alexa Fluor 594 | Thermo Fisher | A-21207 | 1:200 for immunofluorescence |
| Goat anti-rabbit IgG, Atto 647N | Sigma-Aldrich | 40839 | 1:200 for immunofluorescence |

#### Table S2. Oligonucleotides

| # | Oligo name | Sequence 5’-3’ |
| --- | --- | --- |
| 1 | Oligos in EMSA | GAGCATCCGGATCCCCTGGAGAATC |
| 2 | (72 bp) DNA Oligos | GCCACCGGTGGCTTCTTCTAGCCACCGGTGGCTTCTTCTAGCCACCGGTGGCTTCTTCTAGCCACCGGTGGC |
| 3 | 18 bp DNA Oligos | CTAGCCACCGGTGGCTTC |
| 4 | 36 bp DNA Oligos | GCCACCGGTGGCTTCTTCTAGCCACCGGTGGCTTCT |
| 5 | 54 bp DNA Oligos | GCCACCGGTGGCTTCTTCTAGCCACCGGTGGCTTCTTCTAGCCACCGGTGGCTT |
